# Degradation of mucin *O*-glycans by a human gut symbiont requires a complex enzyme repertoire and promotes colonization

**DOI:** 10.64898/2026.03.26.714468

**Authors:** Sadie R. Schaus, Chunsheng Jin, Grete Raba, Gabriel Vasconcelos Pereira, Rajneesh Bains, Carmen R. Cori, Maria-Jose García-Bonete, Marcus Nilsson, Naba D. Salman, Nicholas A. Pudlo, Qinnan Yang, Jining Liu, Jan Holgersson, Stephen G. Withers, Rachel Hevey, Eric C. Martens, Ana S. Luis

**Affiliations:** Department of Microbiology and Immunology, University of Michigan Medical School, Ann Arbor, Michigan, United States; Department of Medical Biochemistry and Cell Biology, Institute of Biomedicine, Sahlgrenska Academy, University of Gothenburg, Gothenburg, Sweden; Proteomics Core Facility at Sahlgrenska Academy, University of Gothenburg, 40530, Gothenburg, Sweden; Department of Chemistry and Biotechnology, Tallinn University of Technology, 12618 Tallinn, Estonia; Department of Chemistry, University of British Columbia, 2036 Main Mall, Vancouver, British Columbia, Canada; Departement of Pharmaceutical Sciences, University of Basel, Switzerland; Department of Biochemistry & Molecular Biology, University of Alicante, Spain; Department of Laboratory Medicine, Institute of Biomedicine, Sahlgrenska Academy, University of Gothenburg, Gothenburg, Sweden; Department of Clinical Immunology and Transfusion Medicine, Sahlgrenska University Hospital, Gothenburg, Sweden; SciLifeLab, University of Gothenburg, Gothenburg, Sweden

**Keywords:** Mucins, colonic *O*-glycans, gut microbiota, mucin-active enzymes, *O*-glycans degradation, gut colonization

## Abstract

Secreted mucins are the major component of the mucus layer that protects intestinal epithelial surfaces by blocking excessive interactions with the microbiota. Mucins are complex glycoproteins decorated with over 100 different *O*-glycans. Some bacteria can utilize mucins and excessive degradation has been associated with disruption of the mucus barrier and inflammation. Despite the importance of mucins, a detailed enzymatic pathway by which gut bacteria degrade colonic mucin *O*-glycans and the impact of this process on gut colonization are unknown. Here, we identified >100 genes that are expressed by the symbiont *Bacteroides thetaiotaomicron* during growth on different *O*-glycan substrates, revealing effects of glycan structure on gene expression. The characterization of 33 glycoside hydrolase enzymes revealed the pathway for colonic *O*-glycan degradation by this bacterium. *In vivo* competition experiments show that multiple exo-acting enzymes targeting mucin capping structures are central to gut colonization and may provide targets to inhibit bacterial mucin degradation.

## Introduction

The 100 trillion intestinal microbes that live in symbiosis with humans ordinarily pose little threat. However, the presence of a dense microbial community (microbiota) just 10-100s of microns away from host tissue presents a challenge to maintaining homeostasis. When microbes or their antigens reach too close to the intestinal epithelium, it may lead to inflammation. Secreted mucus is one of the first layers of defense that protect the epithelium from the microbiota^1^. In the colon, secreted mucus forms two layers. The inner layer is almost devoid of bacteria while the loosely arranged outer layer is heavily colonized by the colonic microbiota^2,3^. Previous studies suggest that the activity of gut bacteria can lead to alterations of the protective mucus layer properties^4^ and diseases such as inflammatory bowel diseases (IBDs)^5,6^, colorectal cancer^7^, pathogen susceptibility^8^ and graft-versus-host disease^9^.

The mucus that covers and protects the colonic epithelium is produced by goblet cells and is mostly composed of secreted mucins. The composition of mucins varies along the gastrointestinal tract with mucin-2 (MUC2) and mucin-5AC (MUC5AC) being expressed in intestine and stomach, respectively^10^. These mucins have proline-threonine-serine (PTS) domains that are heavily decorated with hundreds of *O*-linked glycans attached to Thr or Ser. *O*-Glycans contain D-galactose (Gal), L-fucose (Fuc), *N*-acetyl-D-glucosamine (GlcNAc), *N*-acetyl-D-galactosamine (GalNAc) and sialic acid such as *N*-acetylneuraminic acid (Neu5Ac) and, in most mammals, *N*-glycolylneuraminic acid (Neu5Gc). All mucin-type *O*-glycans originate from a GalNAc residue attached to Ser or Thr, which can be further extended into one of four major core structures. The key difference among cores 1–5 lies in the sugar composition and linkage pattern: Cores 1, 3 and 5 are linear, whereas Cores 2 and 4 are branched, with Core 1/2 built on Gal-β1-3-GalNAc, Core 3/4 on GlcNAc-β1-3-GalNAc and core 5 on GalNac-α1-3-GalNAc backbones. *O*-Glycans also harbor internal sulfate modifications and can be capped at their non-reducing end by blood group epitopes (ABO and Lewis antigens), terminal sulfation and acetylation in sialic acid^10^ (**Figure 1A**). Additionally, the sugar composition, linkages and sizes of *O*-glycan chains are variable along the length of the intestine and between species^11,12^.

**Figure 1.**
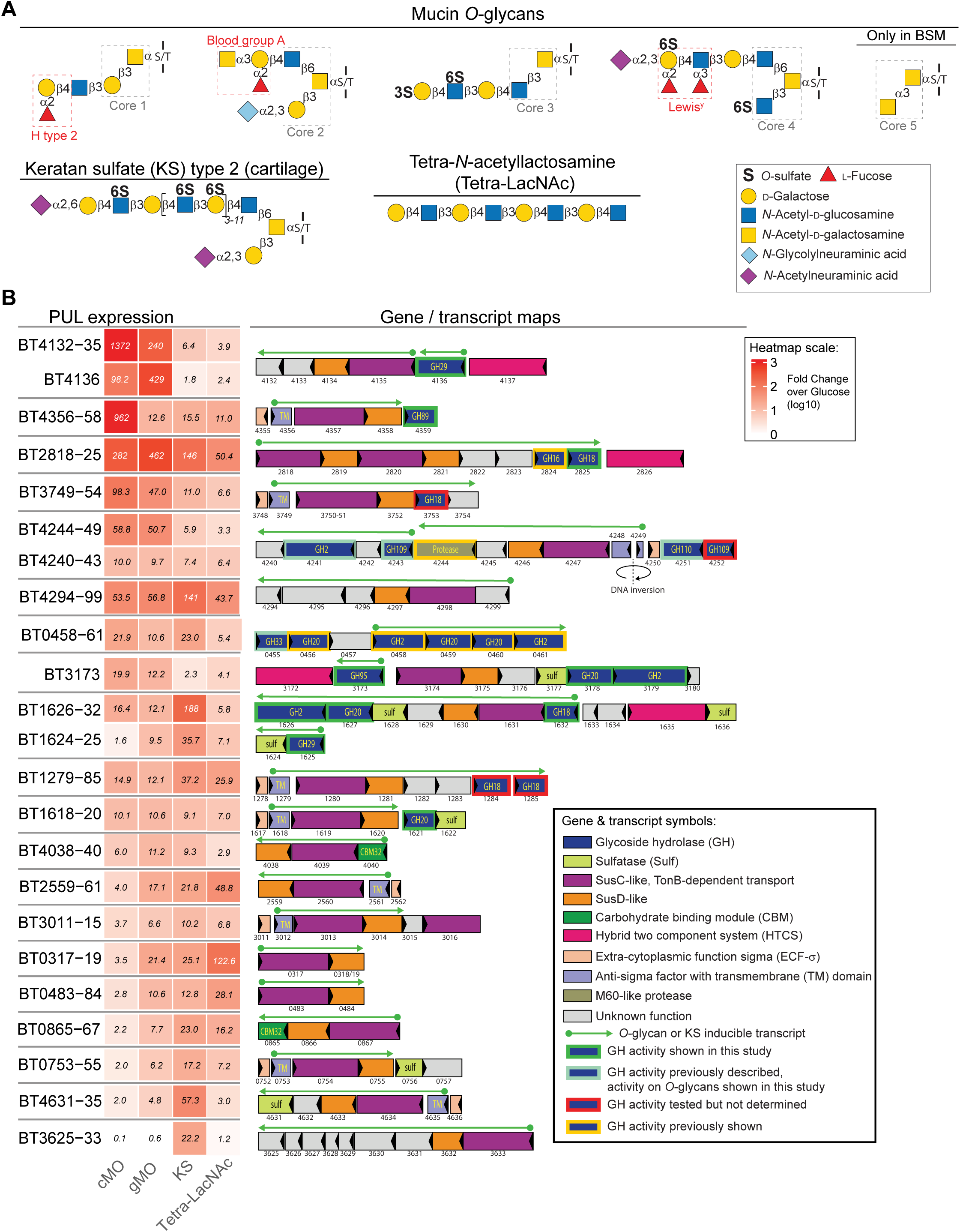
*B. theta* upregulates multiple PULs when grown on different host glycans. (A) Schematic representation of mucin *O*-glycans and closely related substrates used in this study. Core structures and terminal epitopes are highlighted in boxes. Monosaccharide symbols follow the Symbol Nomenclature for Glycans^83^. (B) Heat map of PUL induction by various host glycans. Gene numbers are shown on the left; each box represents the average fold-change (relative to MM-glucose) of all genes within the indicated operon. PUL schematics are shown to the right, with genes detected in RNA-seq data highlighted by green arrows. Only PULs with average fold-change values ≥10 are displayed. BSM, Bovine submaxillary mucin; cMO, porcine colonic mucin *O*-glycans; gMO, porcine gastric mucin *O*-glycans; KS, keratan sulfate; Tetra-LacNAc, tetra-*N*-acetyllactosamine.

The complexity and extensive glycosylation of mucins protect these glycoproteins from bacterial degradation. Indeed, only certain gut bacteria can degrade mucins or its glycans and use it as a source of nutrients, with examples known for members of most major phyla commonly found in the gut^13–17^. Bacteria that forage on mucin or its components need to overcome the chemical complexity of mucin glycans and have likely adapted strategies to both sense structural variations in mucin glycans and express appropriate degradative enzymes. While many bacterial genes and corresponding enzymes involved in this process have been defined^18–23^, the activity of such enzymes on complex *O*-glycans remains unclear and a model for bacterial enzymatic degradation of colonic *O*-glycans remains unknown.

To degrade complex glycans, gut bacteria encode multiple carbohydrate active enzymes (CAZymes) such as glycoside hydrolases (GHs) and sulfatases. GHs are classified in families (GHXX, where XX represents the family number) in the CAZy database based on sequence similarity, which generally reflects conservation of protein fold^24^. The catalytic mechanism and machinery are typically conserved within a GH family; however, exceptions have been reported^25,26^. However, the substrate specificity is not conserved between members of a single family^24^. Therefore, the characterization of *O*-glycan active enzymes is important to expand our limited understanding of how these complex host glycans are degraded and utilized by bacteria.

Two major mucin degraders, *Bacteroides thetaiotaomicron* (*B. theta*) and *Akkermansia muciniphila*, reside predominantly in the outer mucus layer of the colon^27^. Both species can metabolize sialylated and fucosylated *O*-glycans; however, only *B. theta* efficiently grow on sulfated colonic *O*-glycans^28^. Other mucin specialists display narrower substrate preferences. *Ruminococcus gnavus* primarily targets sialylated core 1 structures^29^, whereas *Ruminococcus torques* prefers core-type glycans lacking terminal modifications^17^ and *Bifidobacterium bifidum* predominantly degrades neutral core 1 and core 2 *O*-glycans^30^. Despite several bacteria being able to utilize *O*-glycans, it remains unclear how mucin-degrading species respond to variations in *O*-glycan structures, what are the specific enzymatic mechanisms of degradation of colonic *O*-glycans by bacterium and, which critical ‘early’ steps are necessary to initiate this utilization. To address these knowledge gaps, we investigated the gene regulation and enzymatic mechanisms by which the human symbiont *B. theta* degrades highly sulfated colonic mucin *O*-glycans. Utilization of mucin by this species is emerging as an important contributor to conditions like graft-versus-host disease after irradiation and stem cell transplant^9^ and worsening inflammation in animal models of IBD^6,31^. Our results reveal that growth on different *O*-glycan sources from the stomach and colon activates expression of a partially unique set of gene clusters encoding degradative enzymes. Biochemical characterization of 33 recombinant enzymes reveals the specific activity of these enzymes on complex *O*-glycans and a nearly complete enzymatic pathway by which *B. theta* degrades colonic *O*-glycans. The corresponding deletion of specific enzymatic steps shows that certain steps play strong roles in promoting the ability of *B. theta* to utilize *O*-glycans *in vitro* and to competitively colonize the gut. Overall, this work illuminates the possibility to design specific pharmaceuticals that block key steps in mucin utilization. Such drugs could be used in the future to reduce the deleterious effects of bacterial mucin degradation and its implication in diseases.

## Results

### Variations in mucin O-linked glycan structures activate unique gene expression patterns

The *O*-glycan structures attached to secreted mucins vary in composition along the length of the intestine. In humans and pigs, this variation occurs as increased terminal fucosylation in the stomach and proximal intestine and increased sulfation and sialylation in the colon^12,32–34^. Additionally, gastric and colonic mucins have a higher abundance of core 2 and core 4 structures, respectively, and gastric mucins are decorated with terminal α-GlcNAc epitopes^28,34,35^. We previously determined the global transcriptional responses of *B. theta* during growth *in vitro* with porcine gastric mucin *O*-glycans (gMO) as the sole carbon source^14^. This bacterium typically coordinates expression of multiple carbohydrate utilization genes within clusters termed polysaccharide utilization loci (PULs), which is a common strategy throughout the Bacteroidetes phylum^14,36^. Previous growth on gMO resulted in activation of 82 genes (≥10-fold) that belong to at least 12 different PULs^14^. This is a much larger number of genes than are required to degrade simpler polysaccharides and undoubtedly reflects the complexity of *O*-glycans attached to gastric mucins. However, due to the lack of available source of colonic mucins, the responses of *B. theta* to highly sulfated colonic mucin *O*-glycans, which are more relevant to its colonic habitat, have not been previously measured.

To close this gap, we used a custom preparation of porcine colonic mucin *O*-glycans (cMO) and performed RNAseq-based transcriptional profiling to determine which genes are upregulated during cMO degradation, comparing these responses to growth on gMO (since previous experiments used GeneChip microarrays, gMO transcript analysis was repeated here using RNAseq). As an additional comparison, we also performed transcriptional profiling on keratan sulfate type II (KS), a substrate with a sulfated *N*-acetyl-D-lactosamine (LacNAc) backbone that resembles *O*-glycans (**Figure 1A**). *B. theta* exhibits robust growth on cMO, gMO and KS (**Figure S1A**). Compared to growth on glucose, these substrates activated expression of 370 different genes ≥10-fold in one or more of the conditions tested. Notably, 202 (56%) of these genes belong to a total of 43 different PULs, or close variants of classically defined PULs^14,37^. There was substantial partitioning of responses to a subset of substrates as only 51 (25%) of the activated PUL genes were common to all three substrates (**Figure S1B**). This suggests that *B. theta* has evolved to activate PUL transcription in response to structural diversity of *O*-glycans in different parts of the gastrointestinal tract.

To more deeply probe the responses of individual PULs to variations in the three substrates, we examined the PULs containing genes activated by gMO, cMO and/or KS, adding back any genes that fell below our initial 10-fold expression cutoff. This allowed us to calculate a more accurate fold-change response for each PUL operon and stratify these responses based on the substrates that induce the strongest transcriptional response. Several individual PULs that exhibited transcriptional responses on the substrates tested have previously been associated with degradation of dietary carbohydrates (*e.g*., fructan, ribose, arabinan, arabinogalactan, α-mannan, homogalacturonan) or other host glycans such as *N*-glycans and glycosaminoglycans (chondroitin and heparan sulfatase). Some of these responses are likely due to contaminating fiber polysaccharides and additional host glycans in the colonic mucin preparation (**Table S1**). Because these PULs are unlikely to play direct roles in *O*-glycan degradation they were omitted from further analyses, yielding a list of 20 candidate *O*-glycan PULs (**Figure 1B**).

PULs typically encode degradative enzymes and SusC-like transporters. In total, the activated PUL transcripts that were implicated in *O*-glycan degradation encompass a total of 21 putative degradative enzymes: 17 glycoside hydrolase (GH), 3 sulfatases, and one known M60-like mucin peptidase^38^. However, several PULs harbored genes encoding potential *O*-glycan degrading enzymes (8 GHs and 5 sulfatases) that did not show fold-change differences relative to glucose (**Figure 1B**). Additionally, 4 orphan genes encoding putative enzymes (BT0438, BT3868, BT4050 and BT4394) were also activated in KS and cMO (**Table S1**). Notably, the identified PULs contain a total of 20 SusC-like transporters, which are always encoded upstream of a gene encoding a partner SusD protein with which it complexes during glycan transport^39^, as well as 13 transcriptional sensor/regulators from the hybrid two-component system (HTCS) and extra-cytoplasmic function sigma (ECF-σ) families (**Figure 1B**). This number of transport and sensory functions is far greater than the number involved in *B. theta’s* response to even the most complex dietary fiber polysaccharide, rhamnogalacturonan II^40^, likely due to the additional complexity introduced by the diverse library of *O*-glycans attached to either gastric or colonic mucin combined with the structural variation in the two mucins and KS. HTCS and ECF-σ regulators function by either binding glycans directly in the periplasm^41^ or during transport across the outer membrane^42^, respectively. This suggests that the genes associated with these regulators are likely to be locally activated in response to individual glycan cues. Thus, *O*-glycan linkage, size and decorations such as sulfation, fucosylation and sialylation influence how gut bacteria sense these nutrients.

To test if any *O*-glycan responsive PULs are activated in response to the LacNAc backbone motif common to *O*-glycans and KS, we performed an additional RNAseq experiment using the octasaccharide (Tetra-LacNAc). Interestingly, most *O*-glycan or KS responsive PULs were not as strongly responsive to Tetra-LacNAc as they were to the more complex substrates, suggesting that additional structural features through variations in linkage, branching, or substitution with fucose, sulfate, or sialic acid are required to trigger the glycan sensors associated with these PULs (**Figure 1B**). However, three additional PULs, (BT2559-62, BT0317-19 and BT0483-84) exhibited stronger responses to Tetra-LacNAc than any of the other glycans tested, albeit with variable amounts of induction (28.1-122.6-fold).

All three of these clusters include only genes encoding SusC/D transporter pairs or a transporter pair associated with an ECF-σ regulator (**Figure 1B**). This observation raises the possibility that these systems exist to scavenge non-substituted poly-*N*-acetyl-D-lactosamine (poly-LacNAc) oligosaccharides that are generated by the activity of other enzymes that remove attached fucose, sialic acid, and sulfate modifications.

Previous studies have reported the activity of eight *B. theta* GHs that are encoded in the activated PULs ^23,43–46^. However, it remains unclear if these enzymes are active on complex *O*-glycans. In addition, activities of other putative enzymes encoded in the activated PULs have never been reported. To gain insight into how *B. theta* degrades mucin *O*-glycans, we cloned and purified recombinant forms of the 44 putative glycoside hydrolases (including the 17 GHs encoded in activated PULs) (**Figure S2**) and determined their ability to cleave various oligosaccharides and complex *O*-glycans (**Table S2**).

### Novel endo-acting β-N-acetylglucosaminidases provide early catalytic steps

*Bacteroides* often encode endo-acting enzymes that cleave longer polysaccharides into oligosaccharides that can be imported into the periplasm and further degraded^36^. Recently, BT2824, a glycoside hydrolase family 16 (GH16) was identified as the first *O*-glycan endo-acting β-galactosidase enzyme^47^ (**Figure 1B**). Our transcriptional analysis revealed that four GH18 enzymes were encoded in the activated PULs. BT2825 is found immediately adjacent to the previously characterized endo-acting GH16, while BT1632 is encoded in a separate PUL encoding multiple enzymes, including BT1636 (sulfatase) that is particularly important for degradation of sulfated colonic *O*-glycans and gut colonization^28^. BT1284 and BT1285 are encoded in a PUL without previously characterized enzymes. (**Figure 1B**). To date, GH18 members have not been associated with *O*-glycan degradation. This family is instead known to include endo-acting β*-N*-acetyl-glucosaminidases that cleave GlcNAc-β1,4-GlcNAc linkages found in chitin or *N*-glycans^24^. Phylogenetic analysis of GH18 enzymes revealed that BT1284 and BT1285 are likely endo-β-*N*-acetyl-glucosaminidases. However, BT2825 and BT1632 cluster separately from previously characterized enzymes (**Figure S3A**), suggesting that these enzymes can target a different linkage. BT1632 is also related to another *B. theta* GH18 member (BT3050, **Figure S3A**) that is neither contained in a PUL nor induced by *O*-glycans. Based on these observations, we hypothesized that BT2825, BT1632 and BT3050 target the *O*-glycan poly-LacNAc backbone (**Figure 1A**). To test this, we purified recombinant forms of these GH18 enzymes and tested their ability to cleave a range of LacNAc-substrates that vary in their length and complexity. Analysis of the reaction products showed that these enzymes are indeed endo-acting and display unique substrate specificities (**Table S2**). Consistent with the activity of GH18 family members on β−GlcNAc-linkages, all enzymes cleaved at adjacent sites in the poly-LacNAc chains, generating products with Gal at the non-reducing end (**Figure 2A, B**).

**Figure 2.**
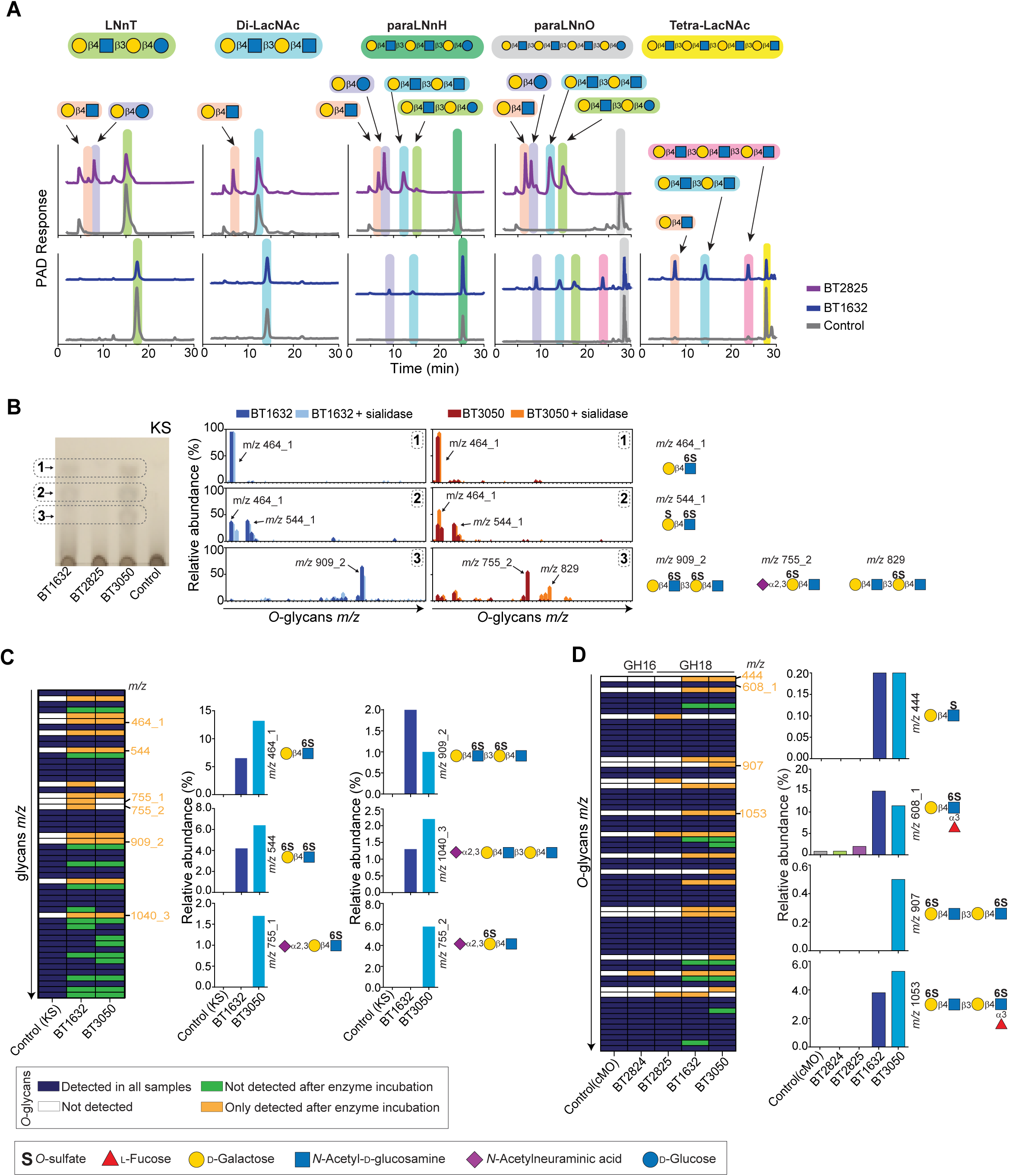
*B. theta* endo-acting enzymes display different activities against host glycan backbones. (A) HPAEC-PAD analysis of reaction products generated by endo-acting enzymes on different oligosaccharides. Initial substrates and products are highlighted in different colors. (B) Thin-layer chromatography of KS digestion products (left panel). Major product bands (1– 3), highlighted in boxes, were analyzed by mass spectrometry before and after incubation with 1 µM sialidase (BT0455) for 16 h at 37 °C (right panel). The most abundant glycans are indicated, with schematic representations shown on the right. (C, D) Representation of glycans detected by mass spectrometry in KS (C) and cMO (D) before and after enzyme treatment (left panels). Relative abundance and corresponding structures for specific *m/z* values are shown on the right. Ions with an underscore followed by a number denote glycans of identical mass but different structures. In all reactions, Control refers to assays performed without endo-acting enzymes under the same conditions. LNnT, lacto-*N*-neotetraose; Di-LacNAc, Di-*N*-acetyl-D-lactosamine; paraLNnH, para-lacto-*N*- neohexaose; paraLNnO, para-lacto-*N*-neooctaose; KS, keratan sulfate; cMO, porcine colonic mucin *O*-glycans.

BT2825 was the only enzyme that acted exclusively on non-substituted poly-LacNAc, displaying a preference for hexasaccharide substrates or longer but not tolerating sulfation or the presence of additional substitutions (**Figures 2A, B**). The activity of BT2825 on tetrasaccharides was dependent on the non-reducing end Gal linkage, with this enzyme only cleaving type 2 glycans (Gal-β1,4-GlcNAc-) (**Table S2**). Although BT1632 and BT3050 are phylogenetically related, their 23% sequence identity (**Figure S2**) suggests possible differences in specificity. Indeed, BT1632 had weak activity against longer poly-LacNAc (≥ hexaoligosaccharides) but displayed a marked activity against KS, whereas BT3050 was only active on KS (**Figures 2A, 2B, S3B and Table S3A)**. Analysis of the main products generated by BT1632 and BT3050 on KS revealed that these enzymes release sulfated oligosaccharides (**Table S3B, C),** suggesting that they recognize sulfate groups within a KS backbone. Both enzymes generated the same major mono- and di-sulfated LacNAc products (band 1 and 2, respectively, in **Figure 2B**), with structures confirmed by enzymatic digestion (**Figure S3C)**. However, BT3050 additionally released sialylated trisaccharide (product 3 in **Figure 2B**), as confirmed by subsequent cleavage with a sialidase (**Figure 2B, S3C**). This suggests that only BT3050 can accommodate sialic acid in the -3 subsite. Indeed, BT3050 was able to cleave synthetic Neu5Ac-α2,3-6SLacNAc-4-nitrophenyl (pNP) but showed almost no activity on a substrate lacking sialic acid (6SLacNAc-pNP) (**Table S4**). The lack of BT3050 activity on sialylated synthetic core 1 (Neu5Ac-α2,3/6-Gal-β1,4-(6S)GlcNAc-β1,3-Gal-β1,3-GalNAc-αC_2_N_3_) also indicates that this enzyme prefers LacNAc in positive subsites (**Table S2**). The mass spectrometry (MS) analysis of the complete activity profile of BT1632 and BT3050 in KS confirmed that the major products are sulfated di- and tetrasaccharides with BT3050 being the only enzyme to release a sialylated trisaccharide. However, the detection of a sialylated tetrasaccharide after BT1632 incubation indicates that this enzyme can act on long sialylated *O*-glycans but cannot tolerate sialic acid in the -3 subsite (**Figure 2C**). To determine if these enzymes are able to cleave cMO we incubated the three GH18s with this substrate, along with the previously characterized GH16 (BT2824)^47^, and determined changes to this complex mixture of glycans by MS (**Figure 2D and Table S3D**). Only BT1632 and BT3050 displayed a clear activity on cMO (**Figure 2D, Table S3D**). Consistent with the results in KS, both enzymes generate oligosaccharides with a sulfate group on the reducing end GlcNAc. Interestingly, the major products also contain fucose at the reducing end (**Figure 2D**), suggesting that BT1632 and BT3050 tolerate fucosylated *O*-glycans. BT2825 did not release any products, consistent with the lack of activity of this enzyme on substituted substrates (**Figure 2B and 2D**). Unexpectedly, the characterized GH16 BT2824 could not cleave cMO (**Table S3D,** note that none of the detected glycans has Gal at the reducing end). Previous studies indicate that BT2824 activity requires the cleavage of sialic acid^47^, which is abundant cMO. It is also possible that BT2824 can cleave longer and less substituted small intestine *O*-glycans but is unable to access the highly substituted cMO. Overall, these results indicate that *B. theta* evolved multiple enzymes with different substrate specificities to generate *O*-glycan oligosaccharides. These different mechanisms might enable the uptake of glycans in different contexts that are not accessible to species lacking homologs of these GH18 enzymes.

### Removal of terminal sugars is essential for subsequent O-glycan degradation

Mucin *O*-glycans can be capped with blood group antigens, Lewis-type fucose, sialic acid, and sulfate, leading to high structural diversity (**Figure 1A**). We previously demonstrated that after *B. theta* grows on cMO *in vitro*, no oligosaccharides could be detected in the growth media^28^, indicating that this bacterium encodes all enzymes necessary to cleave the different terminal epitopes. Additionally, we have shown that a single sulfatase (BT1636) that targets a terminal epitope is critical for the utilization of colonic *O*-glycans. We hypothesized that additional *B. theta O*-glycan active enzymes may also play a key role in mucin utilization. However, so far, few enzymes have been characterized in this species^23,43–45^ and the described activities of microbiota enzymes on cMO are limited to the characterized sulfatases^28^. To close this gap, we characterized the activities of 33 of *B. theta* enzymes on *O*-glycans (**Table S2**).

Fucose is a terminal epitope α-linked to positions O2 of Gal and O3/O4 of GlcNAc. *B. theta* encodes 14 enzymes belonging to families GH95 and GH29, previously known to cleave α1,2 and α1,2/1,3/1,4-fucose, respectively. To define their substrate specificities, we determined the activity of four GH95s and seven GH29s. We found that all four GH95s and four GH29s were active in various substrates, whereas the GH29 BT3956 was only active on pNP-*a*-Fuc (**Figure 3A and Table S2**). Four of the active enzymes were encoded in PULs upregulated by *O*-glycans and one was encoded outside of a PUL (BT1842) (**Table S1**). Two closely related GH29s (BT1625 and BT4136) were active on α-1,3/1,4-fucose linkages (**Figure 3A, S2, S4A-C, Table S2 and S4**). These enzymes were more active against cMO than gMO (**Figure 3B**), which is consistent with colonic mucins carrying higher levels of Lewis-type α1,3/1,4-fucosylated epitopes compared with gastric mucins. Treatment with BT1625 or BT4136 alone resulted in similar amounts of fucose release as the combination of both enzymes (**Figure 3B**), suggesting that both enzymes display similar substrate specificity. Indeed, both of these GH29s displayed marked activity against cMO, with BT1625 and BT4136 cleaving Lewis^a/b/x/y^ structures (**Figure 3C and S4D, Table S5**). Analysis on defined oligosaccharides revealed that BT1625 and BT4136 can cleave internal fucosylated linkages and tolerate α2,3-sialic acid and Lewis epitopes decorated with Gal-3S-sulfation (3S-Gal) and GlcNAc-6S-sulfation (6S-GlcNAc) (**Figure S4A-C**). While these enzymes were inactive on 6’S-Lewis^a/x^ trisaccharide where sulfation is O6-linked to Gal (6S-Gal) (**Table S2**), they both tolerated this sulfation on complex *O*-glycans and the activity was not affected by the presence of double sulfation on Gal and GlcNAc (**Figure 3C**), suggesting that recognition of longer glycans is required for the enzymes to be active on sulfated glycans. The activity of BT1625 and BT4136 was also independent of the type of core structures (**Figure 3C and S4D**). Overall, our results suggest that these enzymes evolved to accommodate the structural variability of *O*-glycans by targeting α1,3/1,4-fucose linkages in different contexts.

**Figure 3.**
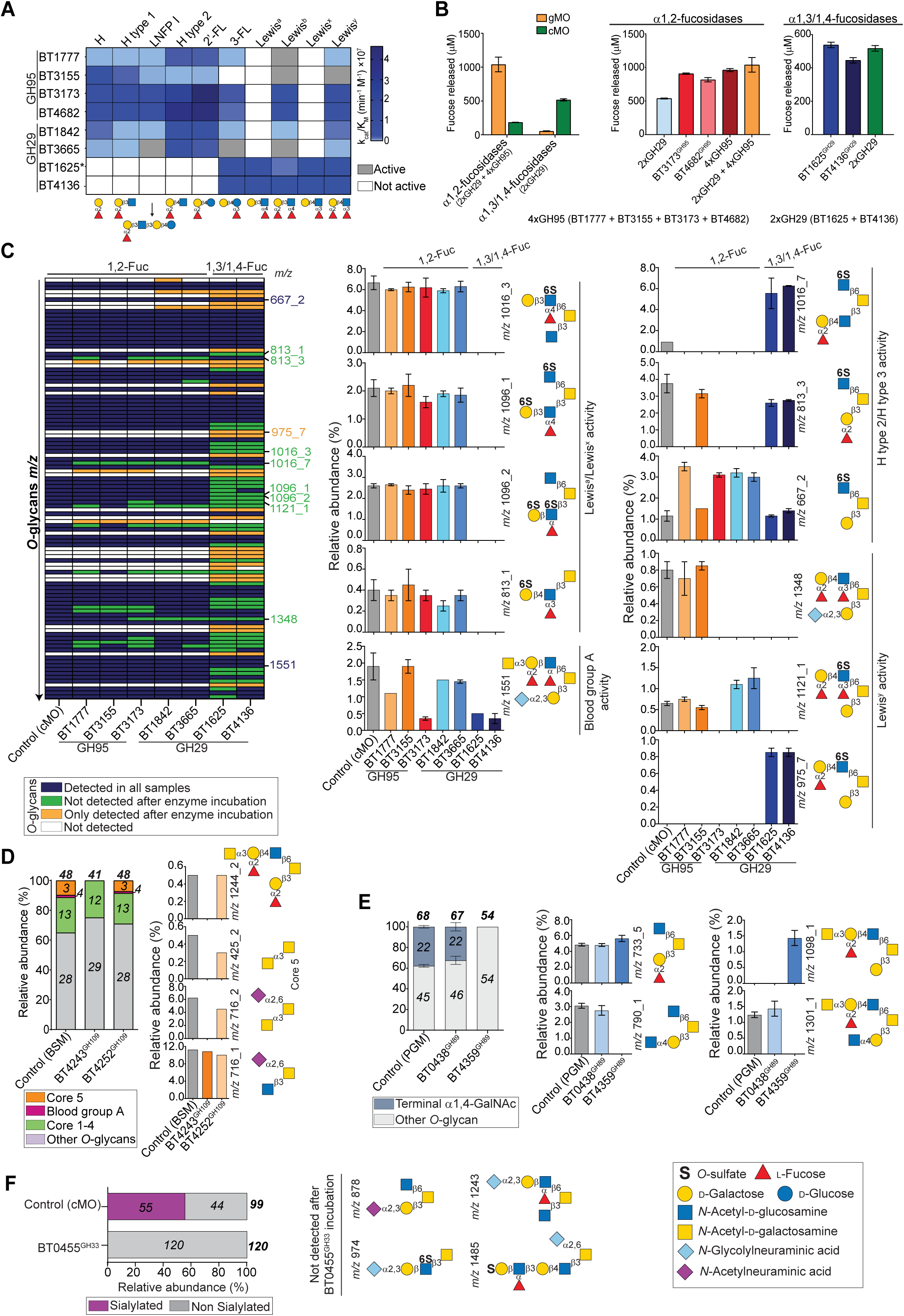
Activity of enzymes cleaving terminal *O*-glycan epitopes. (A) Heatmap of fucosidase activity against various oligosaccharides (**Table S4**). For reactions where it was not possible to determine the enzymatic rate, the dark grey and white colour indicate low activity and no activity, respectively, for reactions set on standard conditions. *BT1625 results previously published^44^. (B) Quantification by HPAEC-PAD of fucose released by different fucosidases or fucosidase combinations (*n*=3-4; error bars are SEM). (C) Glycans detected by mass spectrometry in cMO before (control) and after fucosidase treatment (left). Relative abundance and corresponding structures for specific *m/z* values are shown on the right. Ions with an underscore followed by a number represent glycans with the same mass but different structures (*n* = 2; error bars = SEM). (D) Left: Relative abundance by mass spectrometry of glycan structures detected in BSM (bovine submaxillary mucin) before (control) and after incubation with GH109 enzymes. Numbers indicate the number of glycan structures identified. Right: relative abundance and structures corresponding to specific *m/z* values. (E) Relative abundance of *O*-glycans with or without terminal α1,4-GalNAc detected in porcine gastric mucin (PGM) after incubation with GH89 enzymes (left). Relative abundance and structure corresponding to specific *m/z* values (right panel) (*n* = 3; error bars = SEM). (F) Relative abundance of *O*-glycans with or without terminal sialic acid detected in cMO after sialidase (BT0455) treatment (left). Relative abundance and structure corresponding to specific *m/z* values (right panel). In all reactions, control refers to reactions without enzymes under identical conditions. H, Blood group H; LNFP I, lacto-*N*-fucopentaose I; 2’-FL, 2’-fucosyllactose; 3-FL, 3-fucosyllactose; cMO, porcine colonic mucin *O*-glycans; gMO, porcine gastric mucin *O*- glycans; GalNAc; *N*-acetyl-D-galactosamine

The remaining six fucosidases cleaved α1,2-fucose, although a close analysis of the activities revealed different substrate specificities (**Figure 3A and Table S2**). Consistent with the high abundance of α1,2-fucose reported for gastric mucins, these 1,2-fucosidases released more fucose on gMO than cMO (**Figure 3B**). Two closely related GH95s (BT3173 and BT4682, 74% sequence identity) released almost as much fucose as all six α1,2-fucosidases present together (**Figure 3B and S2**), suggesting that these two enzymes are likely to have similar broad specificities. Indeed, BT3173 and BT4682 displayed very similar activities, targeting H type1/2 and Lewis^b/y^ on oligosaccharides (**Figure 3A, S4B and Table S4)**. In cMO, BT3173 also displayed broad activity and cleaved all α1,2-fucose linkages (**Figure 3C, S4D and Table 5**). BT1842 and BT3665 showed a preference for type 2 linkages in oligosaccharides (**Figure 3A and Table S4**). In cMO, these enzymes were also active on type 3 (Fuc-α1,3-Gal-β1,3-GalNAc) and Lewis^y^, although *O*6-sulfation of GlcNAc (6S-Lewis^y^) blocked the activity (**Figure 3C, S4D and Table S5**). BT3155 displayed activity on H-epitopes but did not tolerate adjacent α1,3-fucose linkages (**Figure 3A and S4B**). Against cMO, this enzyme specifically cleaved type 2 glycans, while no activity was detected in H type 3 or Lewis^y^ (**Figure 3C and S4D**). Using defined oligosaccharides, BT1777 cleaved H epitopes and Lewis^y^ (**Figure 3A and TableS4).** Although, against cMO, this enzyme showed activity in H type 3 and no activity against Lewis^y^ (**Figure 3C, S4D and Table S5**). Overall, these results indicate that *O*-glycan structural variability impacts the activity of α1,2-fucosidases, and that these enzymes have evolved to specifically target different structures within cMO, which could explain why *B. theta* possesses so many of them.

Importantly, the activities of all fucosidases were hindered by the presence of terminal blood A or B groups (**Figure 3C, S4D and Table S2**). *B. theta O*-glycan PULs encode enzymes from families GH109 and GH110 that are active on terminal α-GalNAc (blood group A) and α-Gal (blood group B), respectively. Two GH110 enzymes (BT3160 and BT4251) were previously characterized^45^. Here we confirmed that BT4251 cleaves terminal α-Gal type 1 and type 2 glycans allowing for subsequent α1,2-fucosidase activity (**Figure S4E and Table S2**). The GH109 BT4243 was recently characterized^23^, however, its activity in complex *O*-glycans remains unclear. As described, this enzyme cleaves terminal α1,3-linked GalNAc in all tested oligosaccharides independent of the +1 subsite (**Figure S4F and Table S2**). Consistent with this, in bovine submaxillary mucin (BSM), a substrate decorated with *O*-glycans, BT4243 cleaved the blood group A epitope and core 5 (**Figure 3D and Table S6**), a mucin core previously described in human colonic mucin^48^. Unlikely the previous study^23^, BT4243 activity was not affected by sialylation of the core structure (**Figure 3D**). Two additional GH89s were found active against α1,4-GlcNAc. BT0438 only cleaved pNP-α-GlcNAc (**Table S2)**, while BT4359 displayed activity on gMO, cleaving all terminal α1,4-GlcNAc capping groups detected in this substrate (**Figure 3E, S4G and Table S7**).

The *B. theta* genome encodes a single sialidase (BT0455) that has been previously shown to be active on substrates that mimic the sialic acid linkages found on colonic *O*-glycans^23,43^. Here we show that BT0455 is a broadly active sialidase, which tolerates terminal fucose and sulfate groups and cleaves all sialic linkages found in *O*-glycans. (**Figure 3F, S4H, S4I, Table S2 and S8**). In cMO and released glycans from BSM, the BT0455 sialidase cleaved all Neu5Ac- and Neu5Gc-linkages, and was active against α2,6-linkages attached to the core GalNAc, a major linkage found in human colonic *O*-glycans^49^ (**Figure 3F, S4I and Tables S8A-B**). *B. theta* does not metabolize sialic acid^15,50^. Therefore, the broad specificity of BT0455 sialidase enables the removal of all sialic acid capping structures, providing access to underlying *O*-glycans that can be degraded and utilized by the bacterium. It has been previously shown that acetylation of sialic acid often has an impact on sialidase activity^51^. *B. fragilis* has been shown to encode one enzyme able to remove this acetyl blockage^51^, however, it remains unclear which *B. theta* esterase removes the acetylation on sialic acid. Since colonic mucins are likely highly acetylated further studies to characterize *B. theta* esterases are necessary to understand their role in utilization of colonic mucin. Overall, our detailed characterization of the enzymes targeting terminal epitopes reveals that *B. theta* can overcome the variability present in *O*-glycans by expressing multiple enzymes that target the variety of different linkages found in capping structures.

### Depolymerization of colonic O-glycan backbone and cores is mediated by specific enzymes

After the removal of terminal sulfation, sialylation and fucosylation, we hypothesized that the mucin *O*-glycan backbone and core structures are cleaved by the sequential action of β-galactosidases (GH2) and β-*N*-acetylglucosaminidases (GH20), both of which are abundantly represented in the *B. theta* genome and upregulated PULs (**Figure 1B**). Only two of six GH2s (BT1626 and BT3179) shared high sequence identity (79%) (**Figure S2**), suggesting that the remaining enzymes might target different linkages. BT1626 and BT3179 displayed a preference for type 2 glycans (Gal-β1,4-GlcNAc-linkages) and were also active on lactose (**Figure 4A-B, S5A and Table S9A, S4**). Despite the low activity in type 1 glycans (Gal-β1,3-GlcNAc linkages), both enzymes cleaved β1,3-, β1,4- and β1,6-galactobiose and type 3 glycans (Galβ1,3-GalNAc), when higher enzyme concentrations were used. (**Figure S5A and Table S2**). BT3340 displayed a preference for Gal-β1,4-GlcNAc linkages, although this enzyme was also able to release galactose from type 1 glycans (**Figure 4A-B, S5A and Table S9A**). In contrast, BT4050 and BT4241 preferred Gal-β1,3-GlcNAc linkages (**Figure 4A-B, S5A and Table S9A**). These results are in line with the previously reported activity for BT4241^23^, however, our studies extend the characterization of the substrate specificity to additional epitopes and complex substrates. The presence of fucosylated GlcNAc (Lewis^a/x^) blocked the activity of all galactosidases, but these enzymes tolerated fucosylation in distal positive subsites (LNFP IV) (**Figure S5A and Table S2**). Although GH2s tolerate 6S-GlcNAc, the activity was blocked by the presence of sulfation on the target galactose (**Figure S5B and Table S2**). BT4684 only acted on lactose and β1,4-galactobiose (**Figure 4A, S5A, Table S2 and S4**), suggesting that this enzyme does not tolerate GlcNAc and likely evolved to target human milk oligosaccharides (HMOs) or a different substrate containing these epitopes. These results indicate that *B. theta’s* β-galactosidases can tolerate internal sulfate linkages in GlcNAc, but activity of these enzymes is blocked by the presence of terminal capping structures. Therefore, removal of the terminal substitutions in mucin *O*-glycans is necessary to allow *B. theta* to proceed with their degradation.

**Figure 4.**
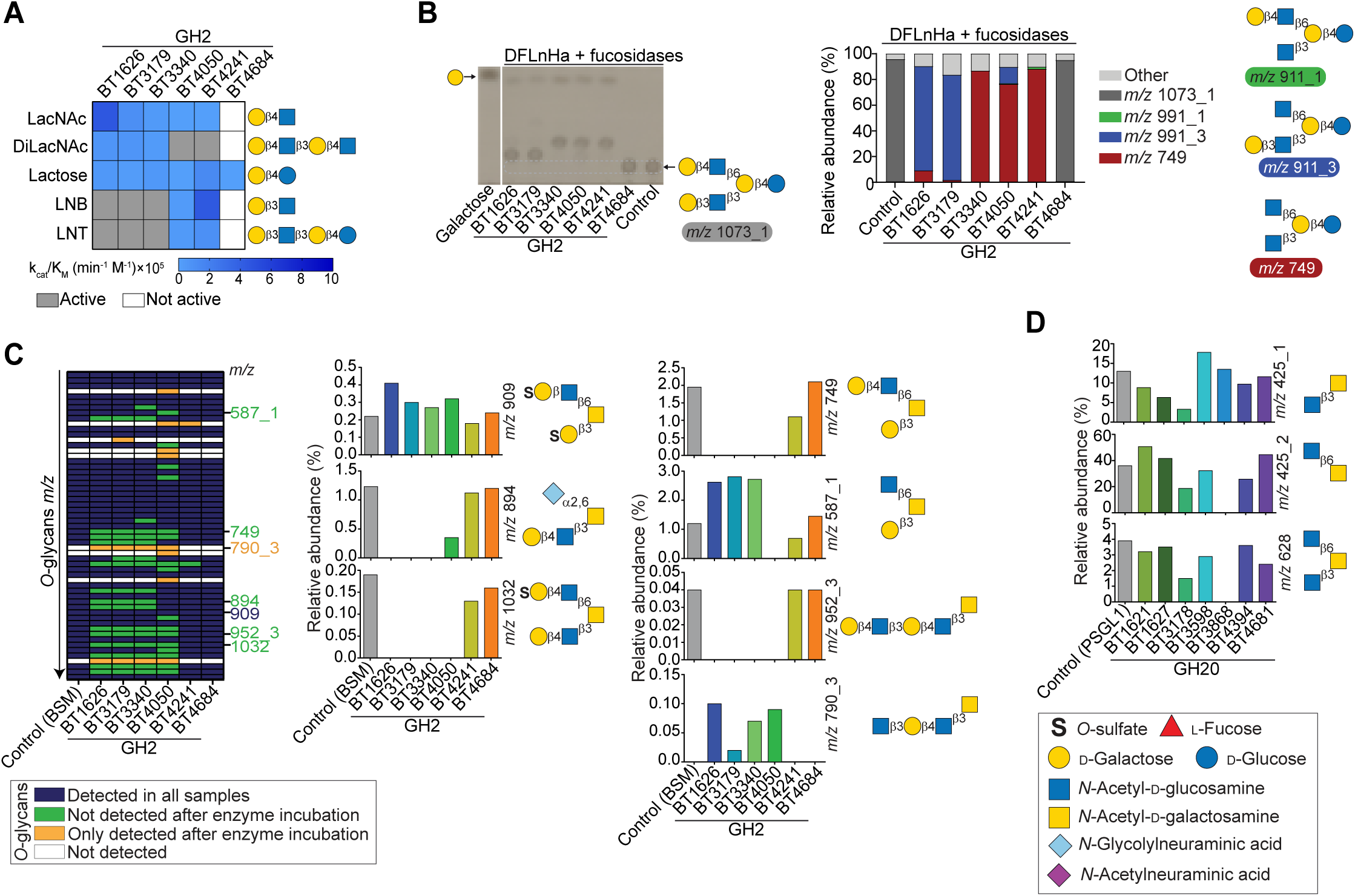
Characterization of *B. theta* GH2 and GH20 enzymes active on *O*-glycans. (A) Heatmap showing the activity of different galactosidases against various oligosaccharides (**Table S4**). For reactions where it was not possible to determine the enzymatic rate, the dark grey and white colour indicate low activity and no activity, respectively, for reactions set on standard conditions. (B) Activity of GH2 enzymes detected by thin-layer chromatography (left panel) and mass spectrometry (right panel). To generate the substrate with m/z 1073_1, difucosyllacto-*N*- hexaose a (DFLnHa) was incubated with 1 μM fucosidases (BT1625 and BT3173) for 16 h at 37 °C. Schematic representations of the structures corresponding to specific *m/z* values are shown on the right. Ions containing an underscore followed by a number represent glycans with identical mass but distinct structures. (C) Glycans detected by mass spectrometry in bovine submaxillary mucin (BSM) after incubation without (control) or with GH2 enzymes (left panel), along with relative abundances and corresponding structures for specific *m/z* values (right panel). To remove terminal epitopes in BSM, all reactions were performed in the presence of 1 μM fucosidases (BT1625 and BT3173) and sialidase (BT0455). Ions containing an underscore followed by a number represent glycans with identical mass but different structures (*n* = 3; error bars = SEM). (D) Relative abundances and corresponding structures for specific *m/z* values of core structures detected on P-selectin glycoprotein ligand-1 (PSGL1) after incubation with GH20 enzymes. To increase the abundance of non-reducing *N*-acetyl-D-glucosamine, PSGL1 was incubated with 1 μM sialidase (BT0455) and galactosidases (BT4241 and BT1626). In all reactions, control refers to reactions without enzymes under identical conditions. LacNAc, *N*-acetyl-D-lactosamine; Di-LacNAc, Di-*N*-acetyl-D-lactosamine; LNB, lacto-*N*-biose; LNT, lacto-*N*-tetraose.

Since the specificity of the β-galactosidases was determined against short oligosaccharides, we next tested their activities against complex *O*-glycans. BSM was used as a model substrate, as it carries shorter *O*-glycans than cMO, allowing us to determine how different cores attached to the protein backbone influence the β-galactosidase activities. Against BSM, with the exceptions of BT4241 and BT4684, all the enzymes were able to cleave terminal galactose residues. Consistent with the results using oligosaccharides, the activity of all β-galactosidases was blocked by the presence of capping sulfation, fucosylation or sialylation (**Figure S5C and Table S9B**). Indeed, pre-treatment of BSM with fucosidases increased the number of *O*-glycans with terminal galactose, thereby resulting in higher β-galactosidase activity (**Figure 4C and Table S9C**). Of note, incubation with sialidase failed to cleave sialic acid from BSM (**Table S9C**), most likely due to the presence of acetylation on sialic acid which blocks the sialidase activity^51,52^. In BSM, the enzymes BT1626, BT3179, BT3340 and BT4050 cleaved galactose from both arms of the core *O*-glycan structures (**Figure 4C and S5C**). Despite the activity of BT4241 on short oligosaccharides, this enzyme showed no activity against Gal-β1,3-GalNAc in BSM. This indicates that BT4241 activity is affected by the presence of the protein backbone. The presence of sialic acid α2,6-linked to core 3 (*m/z 894*) resulted in a decreased activity of BT4050 (**Figure 4C and Table S9C**), suggesting that specific substitutions on *O*-glycans can affect the activity of this β-galactosidase. Although all the active GH2s cleaved synthetic core 2 (**Figure S5A**), when this core is linked to the protein backbone, only BT4050 was active on Gal-β1,3-GalNAc linkage, suggesting specialization for core remnants attached to the polypetide (**Figure 4C, S5C and Tables S9B-C)**. To confirm the activity on core 2, we utilized a glycoprotein (P-selectin glycoprotein ligand-1) decorated with simple core structures. On this substrate, cleavage of the β1,4-Gal linked to core 2 (*m/z 749*) resulted in the accumulation of uncleaved core 2 in reactions containing BT1626, BT3179 and BT3340. However, no core 2 was detected in reactions containing BT4050 (**Figure S5D and Table S9D**), confirming its ability to cleave Gal-β1,3-GalNAc (type 3) linkages attached to glycoproteins. Interestingly, BT4241 showed some activity on Gal-β1,3-GalNAc (*m/z 749*) generating the product *m/z 587_3* with a terminal β1,4Gal that was not cleaved and, that accumulated in the reaction (**Figure S5D and Table S9D**). This indicates that BT4241 can act on type 3 but this activity might be affected by the protein backbone and/or *O*-glycosylation pattern and density. Indeed, it has been previously reported that glycoprotease activities can be dependent on the *O*-glycans displayed in specific mucin backbones^53^. In summary, all six GH2s enzymes function as active β-galactosidase with distinct substrate specificities on mucin O-glycans. BT4684 efficiently cleaves lactose, while BT1626, BT3179 and BT3340 preferentially act on type 2 chain, with the first two displaying broader specificity. In contrast, BT4050 and BT4241 show higher specify for type 1 and 3 chains, with BT4050 also having a limited activity towards type 2 chains.

β-Galactosidases activity on *O*-glycans generates oligosaccharides with a terminal GlcNAc that can be subsequently cleaved by β-*N*-acetylglucosaminidases (GH20). To identify the active GH20 enzymes involved in *O*-glycan degradation, we incubated LNnT with BT1626 (GH2), which removes terminal Gal and generates a trisaccharide with terminal β1,3-GlcNAc. This trisaccharide was then used to screen the activity of the GH20 enzymes, identifying three enzymes (BT1627, BT3868 and BT4681) that fully cleaved the terminal GlcNAc (**Figure S5E**). An additional enzyme, BT4394, displayed weak activity on this substrate, while three additional enzymes (BT1621, BT3178 and BT3598) were inactive. Although, all these enzymes were able to specifically hydrolyze pNP-β-GlcNAc, BT3868 and BT4681 were ∼100-fold more active than BT1621, BT3178 and BT4394 (**Table S4**). Co-incubation of Tetra-LacNAc with three GH20s (BT1627, BT3868 and BT4681) and the β-galactosidase BT1626 led to the complete cleavage of Tetra-LacNAc to monosaccharides (**Figure S5F**), indicating that GH20s and GH2s act sequentially to fully cleave the *O*-glycan backbone. Consistent with previous results, BT4394 was very weakly active, whereas BT1621, BT3178 and BT3598 were inactive. Of note, although BT1627 and BT3178 share 76% sequence identity (**Figure S2**), only BT1627 was active on *O*-glycans. Analysis of the predicted structures revealed that despite BT1627 and BT3178 sharing relatively high sequence identity, multiple residues in close proximity to the active site are not conserved between these enzymes (**Figure S5G**) likely due to having a role in substrate recognition; however, further structural studies are required to investigate the substrate specificity determinants of these GH20 enzymes. Importantly, the differences in activity of these two closely related proteins highlight the relevance of biochemically characterizing CAZymes in order to understand their biological role.

Colonic mucins can be decorated with core structures containing β1,3/β1,6-GlcNAc. Incubation of GH20 enzymes with synthetic core 2 and core 4 oligosaccharides revealed that BT3868 was the only enzyme that could cleave GlcNAc from these structures (**Figure S5H**). It cleaves both β1,3 and β1,6 linkages, and the activity on core 2 indicates that this enzyme tolerates the presence of Gal β1,3-linked to GalNAc, while in core 4, BT3868 displayed a preference for the β1,3-linkage (**Figure S5I and Table S10A**). To test whether the β1,3-branch in core 2/4 blocked the activity of additional GH20s, we incubated these enzymes with the disaccharide GlcNAc-β1,6-GalNAc generated by enzymatic digestion of the β1,3-branch in core 2/4. Of the seven enzymes tested, only BT3868 was able to cleave this linkage (**Figure S5H**). Testing of the enzymes on core structures linked to a protein showed that only BT3868 was active, cleaving GlcNAc-β1,6-GalNAc linear or linked to core 4, but with no activity on GlcNAc-β1,3-GalNAc (core 3) (**Figure 4D and Table S10B-C**). This latter result was surprising since BT3868 displayed a preference for GlcNAcβ1,3GalNAc in the synthetic substrate. This indicates that the presence of a protein backbone is most likely interfering with substrate recognition. Overall, the characterization of *B. theta’s* β-*N*-acetylglucosaminidases revealed that this bacterium has evolved multiple GH20 enzymes that act downstream of the galactosidases to cleave the *O*-glycan backbone into monosaccharides, which can be utilized for growth. Interestingly, only BT3868 displayed both β1,3 and β1,6-*N*-acetylglucosaminidase activities, indicating that this enzyme might have key role on utilization of core structures.

### Depolymerization of diverse colonic O-glycans requires a suite of sequential B. theta enzymes

Biochemical characterization of the *B. theta* enzymes active on *O*-glycans allowed us to identify the substrate specificity of 33 enzymes. We next hypothesized that these enzymes work in sequential order such that enzymes targeting the different glycan capping epitopes act first prior to subsequent cleavage of the *O*-glycan backbone. To test this, we incubated defined synthetic oligosaccharides that harbor the different terminal epitopes with specific enzymes that target these different linkages. Degradation of blood groups A and B requires initial cleavage by the enzymes GH109 (BT4243) and GH110 (BT4251), respectively, which then allow the GH95 enzymes to cleave α1,2-Fuc linkages (**Figure S6A**). To cleave Lewis antigens, the fucosidases need to remove all fucose linkages prior to the activity of the enzymes cleaving the backbone (**Figure 5A and S6B**). Importantly, both GH95 (BT3173 and BT4682) and GH29 (BT1625 and BT4136) have similar activities on Lewis^b/y^ indicating that these enzymes can tolerate fucose linked to the adjacent sugar and there is no preferential order of action (**Table S4**).

**Figure 5.**
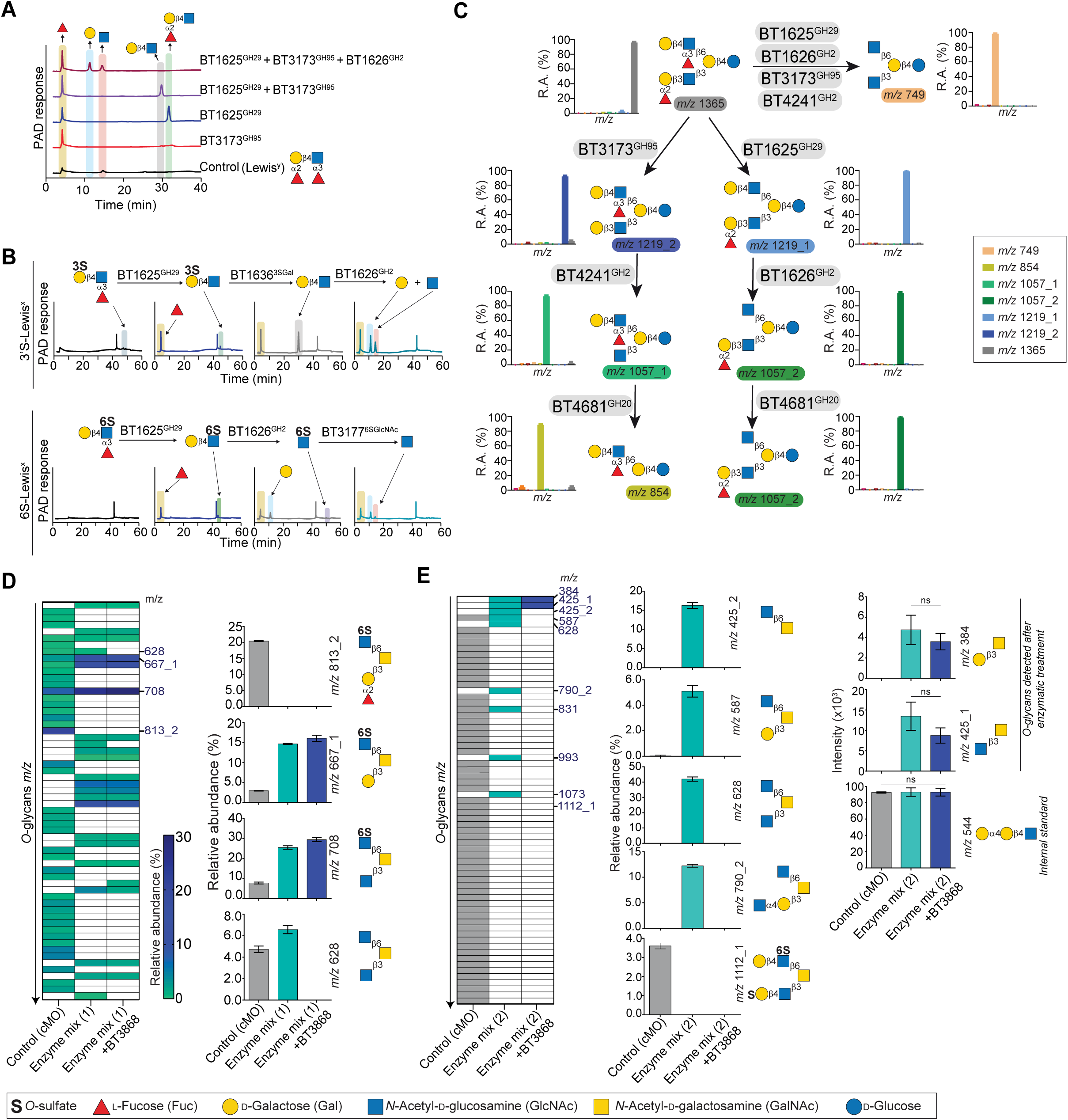
Depolymerization of glycans by *B. theta* enzymes. (A) HPAEC-PAD analysis of sequential degradation of the Lewis^y^ epitope. Products from each enzyme are highlighted in different colours. The initial substrate was not detected in control sample by this technique. (B) Sequential degradation of 3’S-Lewis^x^ (top) and 6S-Lewis^x^ (bottom) and analysis of reaction products (arrows) by HPAEC-PAD. (C) Mass spectrometry of difucosyllacto-*N*-hexaose a depolymerization showing the schematic structures and relative abundances of products. Ions with an underscore and number denote isomeric glycans of identical mass. Reactions were performed under standard conditions with 0.1 mM of substrate. (D, E) Degradation of porcine colonic mucin *O*-glycans with enzyme mixtures. Representation of the glycans detected by mass spectrometry after incubation without (control) or with enzymes and relative abundance and structures for specific *m/z* values. In (E), intensities of the undigested internal control (Gal-α1,4-Gal-β1,4-GlcNAc), core 1 (Gal-β1,3-GalNAc), and core 3 (GlcNAc-β1,3-GalNAc) are also shown (please note that the intensity of internal control remains unchanged). Substrates were incubated with mix 1 or 2 in absence or presence of BT3868. Student’s *t*-test: ns (non-significant) Enzyme mix 1: 1 μM each of BT0455, BT4243, BT1625, BT3173, BT1636, BT1624, BT1626, BT4241, BT1628, BT4681, BT0438. Enzyme mix 2: mix 1 plus 1 μM BT4394. In all reactions, control represents the reactions without enzymes under identical conditions. When possible, the locus tag of each enzyme is followed by the GH family or sulfatase specificity in superscript. R.A., relative abundance; GHXX, glycoside hydrolase family XX; 3SGal, galactose-3*O*-sulfate; 6SGlcNAc, *N*-actetyl-D-glucosamine-6*O*-sulfate

Sulfation strongly impacts *O*-glycan degradation, and the removal of these sulfate groups is required for *B. theta* to utilize colonic *O*-glycans *in vitro* and *in vivo*^28^. We have previously characterized 11 sulfatases that together can target all the sulfated linkages found in *O*-glycans^28^. Here, we show that the presence of sulfate groups that are either 3S-Gal or 6S-GlcNAc did not impact the activity of fucosidases on Lewis^a/x^ (**Figure 5B and S6C**). However, the presence of 6S-Gal on Lewis^a/x^ blocked the activity of all characterized 1,3/1,4-fucosidases (**Table S2**) and, so far, the mechanism by which 6’S-Lewis^a/x^ oligosaccharides are cleaved remains unclear. When Lewis^a/x^ is decorated with 3S-Gal (3’S-Lewis^a/x^), the fucosidases (GH29) generate a disaccharide with a terminal sulfate group that is removed by a 3S-Gal sulfatase (BT1636^28^). The cleavage of this terminal sulfate group is necessary to enable the galactosidase (GH2) activity (**Figure 5B and S6C)**. In the case of 6S-GlcNAc (6S-Lewis^a/x^), upon the removal of fucose, galactosidases cleave the terminal galactose enabling the 6S-GlcNAc sulfatase (BT3177^28^) to remove the sulfate group (**Figure 5B and S6C**).

While the presence of terminal sialic acid blocks the activity of galactosidase (GH2) enzymes, the 1,3/1,4-fucosidases (GH29) are more tolerant and can removed the fucose in sialyl-Lewis^a/x^ (**Figure S6D**). Likewise, the presence of fucose 1,3/1,4-linked to GlcNAc in sialyl-Lewis^a/x^ did not impact the activity of the sialidase (GH33) (**Figure S6D**). The combined activity of the sialidase, fucosidases and sulfatases thus enables the removal of the terminal epitopes allowing the galactosidases and *N*-acetylglucosaminidase to sequentially cleave the *O*-glycans backbone. To validate this concept, we utilized a complex HMO oligosaccharide with H type 1 and Lewis^x^ terminal epitopes (**Figure 5C and Table S11A**). The fucosidase activity was necessary to unblock the sequential degradation of this oligosaccharide, with the 1,2- and 1,3-fucosidases being specifically active on H type 1 and Lewis^x^, respectively. The non-substituted galactose was cleaved by the galactosidases, whereas no activity was detected on the fucosylated branch (*m/z 1057_1/2*). Finally, the GH20 BT4681 only cleaved the GlcNAc-β1,3-Gal linkage (**Figure 5C**), consistent with our observation that BT4681 is active on terminal β1,3-GlcNAc linkages in poly-LacNAc but inactive against core 2, 3 and 4 (**Figure 4D and S5H**), suggesting that this enzyme cannot tolerate β1,6-GlcNAc linkages or GalNAc in +1 subsite (GlcNAc-β1,3-GalNAc).

Enzymatic assays on oligosaccharides indicate that *B. theta* encodes all the enzymes necessary to degrade colonic *O*-glycans. To confirm this, we combined 12 enzymes (mix 1), chosen for their ability to target all *O*-glycans linkages, and incubated this enzyme mix with cMO to test if these glycans were fully degraded. Enzyme mix 1 contained enzymes that cleave sialic acid (BT0455), blood group A (BT4243), α1,2-fucose (BT3173), α1,3/1,4-fucose (BT1625), α-GlcNAc (BT0438), β1,4-Gal (BT1626), β1,3-Gal (BT4241), β1,3-GlcNAc (BT4681) and sulfated linked 3S-Gal (BT1636), 6S-Gal (BT1624) and 6S-GlcNAc (BT1628)^28^. This enzyme mix was incubated with cMO with or without BT3868, the only GH20 enzyme that can cleave GlcNAc from core 2, 3 and 4 (**Figure S6E**). We hypothesized that the enzyme mix lacking BT3868 would degrade the cMO to core structures, while including BT3868 would enable the complete degradation of these glycans. Of 41 *O*-glycans present in cMO, only five remained upon treatment with the enzyme mix, while 18 new *O*-glycans were generated as enzymatic products (**Figure 5D and Table 11B**). This indicated that the majority of the *O*-glycans were indeed degraded by our mix of 11 enzymes. While the presence of BT3868 lead to degradation of core 4 (*m/z 628*), multiple *O*-glycans remained after the enzyme treatment, these being highly decorated with sulfate groups on GlcNAc. Indeed, core 2 and 4 containing 6S-GlcNAc represented more than 50% of the residual *O*-glycans (**Figure 5D and Table 11B**), indicating that the 6S-GlcNAc sulfatase included in our mix failed to remove the “blocking” sulfate groups decorating specific oligosaccharides. To bypass this problem we added GH20 BT4394, which was previously described as an *N*-acetyl-(6-sulfo)-glucosaminidase^46^, to our enzyme cocktail (enzyme mix 2) (**Figure S6E**). The addition of this enzyme mix 2 (without BT3868) resulted in degradation of 58 out of 59 *O*-glycans in cMO (**Figure 5E and Table S11C**). This enzymatic treatment led to the accumulation of seven new glycans with a terminal GlcNAc β-linked to the core. No 6S-GlcNAc glycans remained, suggesting that BT4394 activity may be required to cleave these sulfated epitopes. Adding BT3868 to enzyme mix 2 resulted in the cleavage of all initially detected *O*-glycans in cMO, with only two core glycans remaining (**Figure 5E)**. The detection of core 1 and core 3 is consistent with our previous results showing that BT4241 failed to cleave Gal-β1,3-GalNAc (core1) (**Figure 4C**) and BT3868 did not act on GlcNAc-β1,3-GalNAc (core 3) in complex substrates (**Figure 4D**). Overall, we demonstrate for the first time that cMO can be degraded through a combination of at least 13 enzymes expressed by *B. theta*.

### Key O-glycan catalytic steps are essential for growth in vitro and for in vivo competition

We previously demonstrated that a single sulfatase (BT1636) plays a prominent role in colonic mucin utilization by *B. theta*^28^. Given the variability in mucin *O*-glycan structures, and the sequential mode of action of *O*-glycan active enzymes, we hypothesized that additional enzymes could also be critical for *B. theta* growth on mucin oligosaccharides *in vitro* as well as for colonization *in vivo.* To pinpoint these key *O*-glycan processing catalytic steps, we generated *B. theta* mutants that are diminished in single catalytic activities (*e.g*., all 6 α2-fucosidases) or in multiple activities simultaneously.

Several studies have shown that cell surface endo-acting enzymes are critical for initiating utilization of polysaccharides by *Bacteroides*^44,54–59^. Given the presence of BT1632 at the cell surface (**Figure S7A**) and the ability of the *O*-glycan endo-acting enzymes to cleave longer oligosaccharides into shorter oligosaccharides we hypothesized that the activity of the three GH18s (BT1632, BT2825, BT3050) and the GH16 (BT2814) would be critical for *B. theta* growth on *O*-glycans. We generated a “Δ4× *endo*” mutant lacking the genes encoding these four enzymes. However, the Δ4× *endo* mutant did not display a growth defect compared to the wild-type (WT) strain on gMO, cMO or KS (**Figure 6A**), suggesting that these endo active enzymes are not essential for *in vitro* growth on *O*-glycans, or that additional unidentified enzymes fulfill similar roles.

**Figure 6.**
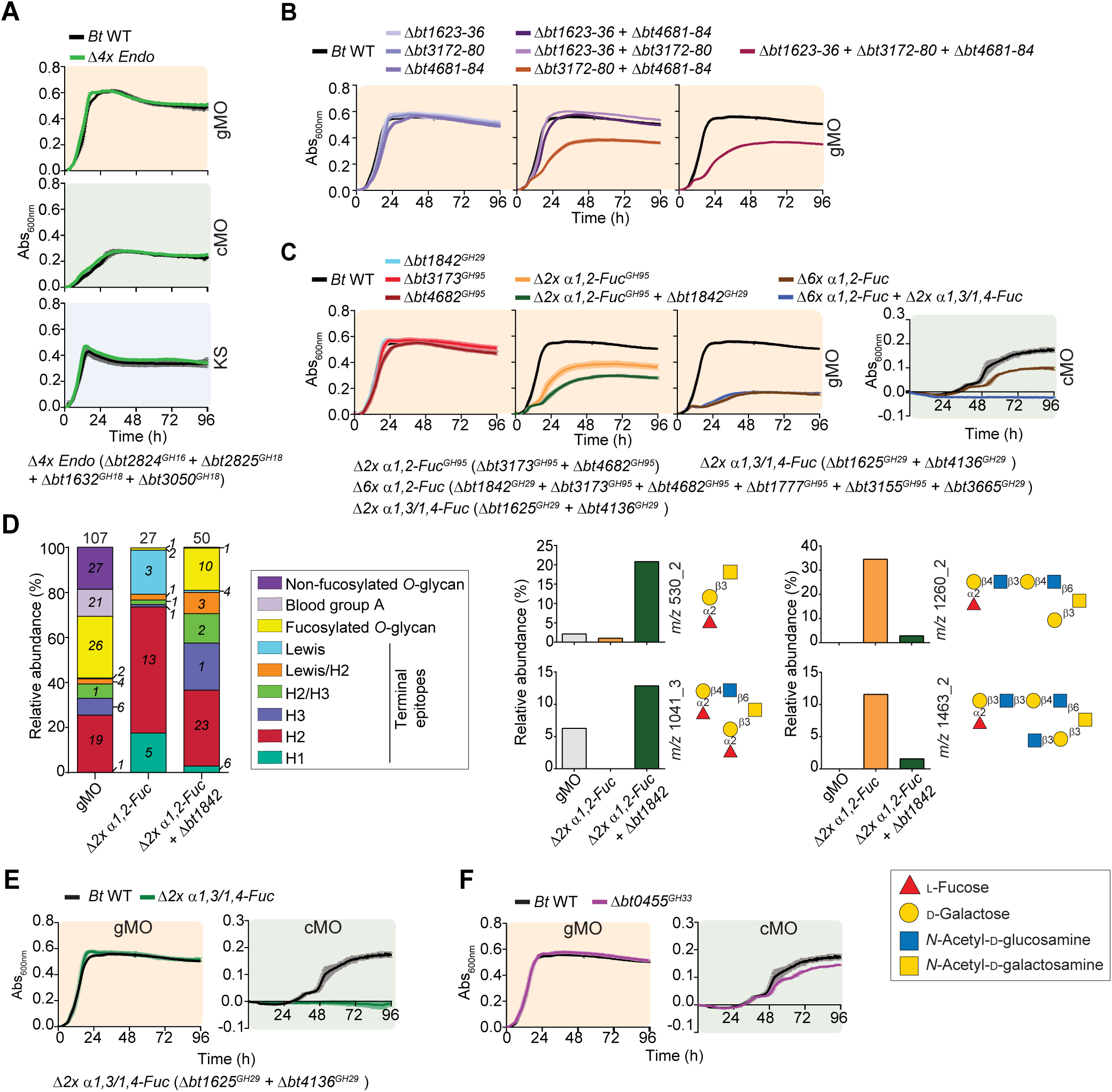
Fucosidases and sialidase activities are required for growth on *O*-glycans. (A–C, E, F) Growth of *B. theta* wild-type (*Bt* WT) and gene-deletion mutants (Δ*btxxxx*) on gMO (10 mg/ml), cMO (5 mg/ml), or KS (5 mg/ml). Lines represent the mean of biological replicates (*n* = 3, except for cMO where *n* = 2; error bars, SEM). To improve clarity, the *Bt* WT growth curve (in black) was added to all panels. (D) Relative abundance of *O*-glycan epitopes detected by mass spectrometry in gastric mucin *O*-glycans (gMO) before and after growth of fucosidase mutants. For four of the most abundant glycans that accumulated in the culture supernatant of the mutants, the corresponding structures and relative abundances across all samples are shown. When applicable, the locus tag of each enzyme is followed by its GH family in superscript. gMO, gastric mucin *O*-glycans; cMO, colonic mucin *O*-glycans; KS, keratan sulfate.

Many of the *B. theta* mucin-degrading enzymes are encoded in PULs (**Figure 1B**). We next investigated the contribution of three PULs that are induced to varying degrees *in vivo* and/or during *in vitro* growth on mucin substrates: *bt1623-36*, *bt3172-80*, and *bt4681-84*. Loss of each individual PUL resulted in growth like WT on gMO. However, the combined deletion of two PULs (Δ*bt3172-80 +* Δ*bt4681-84*) resulted in a growth defect relative to the WT strain, and this phenotype remained unchanged when the third PUL was deleted (Δ*bt1623-36 +* Δ*bt3172-80 +* Δ*bt4681-84*) (**Figure 6B**). Multiple oligosaccharides were detected in the culture supernatant of the triple PUL mutant suggesting that this mutant lost the ability to utilize these glycans (**Figure S7B**). These results suggest that PULs BT3172-80 and BT4681-84 are key in gMO utilization and that cumulative loss of key mucin PULs, but not single PULs, reduces the ability of *B. theta* to metabolize gMO as a nutrient source. Next, we sought to identify the key enzymes in the BT3172-80 and BT4681-84 PULs. These PULs encode two closely related α1,2-fucosidases (BT3173 and BT4682) (**Figure 1B and S2**). Single mutants lacking BT3173 or BT4682 grew similarly to WT on gMO. But a mutant lacking both fucosidases (Δ*bt3173 +* Δ*bt4682,* herein Δ*2*× *α1,2-Fuc*) displayed a growth defect similar to the one observed with the double deletion of PULs Δ*bt3172-80 +* Δ*bt4681-84* (**Figure 6C**). Therefore, the defect observed with the double PUL mutant is likely to be driven by loss of the α1,2-fucosidases, which may be especially important on gMO glycans that frequently terminate in this linkage. Interestingly, the growth defect of the Δ*2*× *α1,2-Fuc* mutant was fully recovered to WT levels when the recombinant enzymes were added to the growth media (**Figure S7C**). The aerobic incubation of the WT or Δ*2*× *α1,2-Fuc* with gMO resulted in release of Fuc by enzymes at the cell surface in WT, whereas the mutant failed to release Fuc (**Figure S7D**). This suggests that at least one of the deleted fucosidases (BT3173 or BT4682) may be located at the cell surface. We generated additional mutants containing single or combinatorial gene deletions of characterized fucosidases. In gMO, the mutants lacking multiple fucosidases demonstrated a gradient of growth, with a triple mutant (Δ*2*× *α1,2-Fuc* + Δ*bt1842*) having a more severe phenotype than the double fucosidase mutant and the most severe phenotype being a strain lacking all six characterized α1,2-fucosidases (Δ*bt3173 +* Δ*bt4682 +* Δ*bt1777 +* Δ*bt3155 +* Δ*bt3665 +* Δ*bt1842,* herein Δ*6*× *α1,2-Fuc*) (**Figure 6C**). Consistent with the growth phenotypes, the accumulation of oligosaccharides in the culture supernatant of the α1,2-fucosidase mutants indicate that these mutants are unable to utilize some glycans (**Figure S7B and S7C**). Indeed, the MS analysis of the glycans in the culture supernatant of Δ*2*× *α1,2-Fuc* mutant revealed that 22 out of the 27 detected oligosaccharides have α1,2-Fuc at their non-reducing end (**Figure 6D and Table 12**). Interestingly, the triple mutant Δ*2*× *α1,2-Fuc* + Δ*bt1842* accumulated additional glycans, with two blood group H type 3 glycans accounting for 30% of the glycans that remained non-utilized in its culture supernatant (**Figure 6D**). This suggests that BT3173 and BT4682 are necessary for the utilization of α1,2-fucosylated *O*-glycans and that BT1842 fucosidase is key to the utilization of H type 3 glycans.

In contrast, loss of the two α1,3/1,4-fucosidases, which target a linkage rarely found in gMO, did not result in a growth defect relative to WT (**Figure 6C**). The additional loss of the two α1,3/1,4-fucosidases in combination with the Δ*6*× *α1,2-Fuc* mutant resulted in a similar growth defect to that observed with the Δ*6*× *α1,2-Fuc* mutant alone (**Figure 6C**), further reinforcing the conclusion that α1,3/1,4-fucosidase activities are dispensable for *B. theta* growth on gMO. In contrast to gMO, on cMO the Δ*6*× *α1,2-Fuc* mutant demonstrated only a moderate growth defect while the α1,3/1,4-fucosidase mutant failed to grow (**Figure 6C and 6E**). Overall, these results indicate that α1,2-fucosidases are critical for utilization of gMO, while α1,3/1,4-fucosidase are needed by *B. theta* to access cMO as a nutrient by *B. theta*. On a different batch of cMO, the Δ*6*× *α1,2-Fuc* mutant displayed the same moderate growth defect, although the phenotype of the α1,3/1,4-fucosidase was not as severe (**Figure S7E)**. Therefore, the phenotypes of these mutants are likely dependent on the abundance of terminal epitopes targeted by the deleted enzymes in each preparation. Overall, these results highlight the need to use colonic substrates in gut microbiota studies aiming to understand how these bacteria interact with mucins.

Sialylation is a common feature in mucin *O*-glycans. To assess the contributions of *B. theta* sialidase, we grew mutant lacking BT0455 on gMO or cMO. In gMO Δ*bt0455* grew similarly to WT (**Figure 6F**) suggesting that sialidase activity is dispensable for growth on this substrate. This is corroborated by MS observations that our gMO preparation harbors relatively few sialylated *O*-glycans (**Table S12**). On cMO, the mutant Δ*bt0455* showed a growth defect compared to the WT strain however the magnitude of this effect was batch-dependent (**Figure 6F and S7E**). This indicates that sialidase activity can have an impact on sialylated cMO utilization which, is in agreement with our enzymatic studies where this enzyme was necessary for the early steps of *O*-glycan depolymerization.

To identify key catalytic activities that play a role *in vivo*, we competed several of the *B. theta* mutants lacking different sets of enzymes against the WT strain in gnotobiotic mice that were fed a fiber-free diet to enhance reliance on mucin *O*-glycans as carbon source^60^. Consistent with its lack of growth defect in *O*-glycans, the Δ4× *endo* mutant did not exhibit an *in vivo* colonization defect in competition with WT, even when competed over 11 weeks (**Figure 7A**), suggesting that these four *endo* enzymes (BT1632, BT2824, BT2825, and BT3050) are dispensable for *B. theta in vivo* colonization. Based on the ability of this mutant to grow on KS, we hypothesize that there are additional endo-acting enzymes encoded in the *B. theta* genome that are functionally redundant with these four endo-acting enzymes. However, the attempted characterization of additional recombinant GH18 enzymes (**Table S2**) or other protein of unknown function did not identify additional activities and such endo-acting enzymes have not yet been identified.

**Figure 7.**
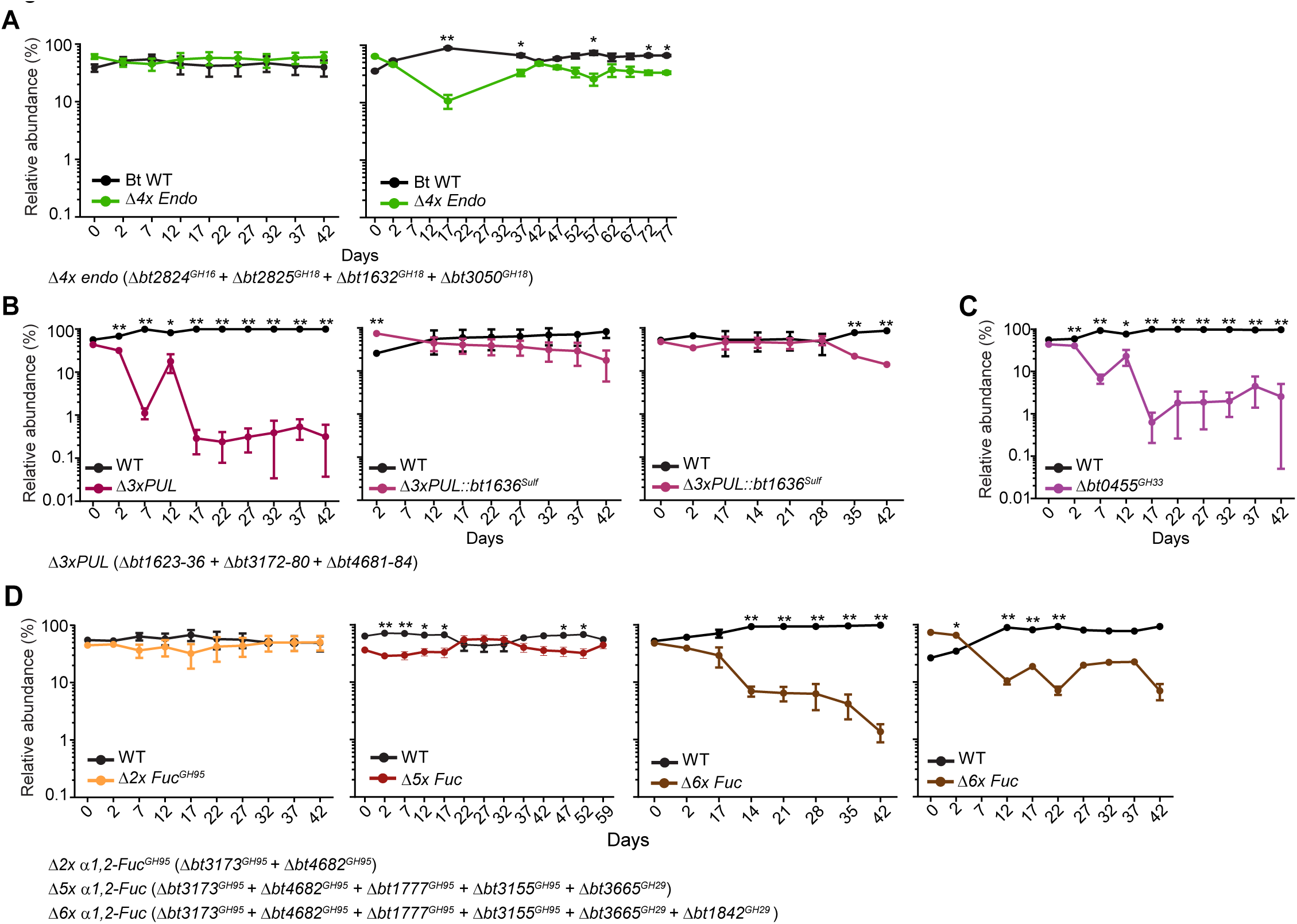
Fucosidases and sialidase are required for competitive fitness *in vivo*. (A–D) Fecal relative abundance during *in vivo* competitions in gnotobiotic mice fed a fiber-free diet and inoculated with *B. theta* wild-type (Bt WT) and mutant strains (Δ*btxxxx*). Relative abundance at time 0 reflects the composition of the gavaged inoculum. Student’s *t*-test: \**p* < 0.05, \*\**p* < 0.01. *n* = 3–8; error bars indicate SEM.

Next, we investigated the impact of PUL deletions on *B. theta in vivo* fitness. The loss of three PULs (Δ*bt1623-36 +* Δ*bt3172-80 +* Δ*bt4681-84*) encoding thirteen enzymes (**Figure 1B**) led to a severe competitive defect (**Figure 7B**). However, this defect was rescued by complementing back only BT1636, a sulfatase that has been previously shown to be important for *in vivo* competition^28^. This reminds us that individual enzymes with unique activity, such as the key sulfatase^28^, may be essential for fitness, while the deletion of enzymes with homologs encoded in the genome may not affect the mutant fitness due to a compensation of the deleted activities by additional enzymes with a high degree of catalytic redundancy.

Finally, we explored the impact of specific *O*-glycan-targeting enzymes on *B. theta in vivo* fitness. Since colonic mucins are also highly sialylated^28^ we tested weather the sialidase activity is important for gut colonization. The *in vivo* competition of Δ*bt0455* against the WT demonstrated a strong competitive defect (**Figure 7C**). Despite the weak phenotype of the sialidase mutant during the *in vitro* growth on cMO (**Figure 6F and S7E**), BT0455 plays a key role in promoting *B. theta* access to mucin glycans *in vivo.* This highlights the importance of complementing *in vitro* results with *in vivo* experiments since the linkages that are most abundant or important could differ between the two. The deletion of two key α1,2-fucosidases (BT3173 and BT4682) did not reduce competitive fitness (**Figure 7D**), despite this mutant exhibiting a defect on gMO *in vitro*. This is also consistent with the lack of a fitness phenotype for the triple PUL mutant, since these fucosidases are encoded within those PULs. Since our *in vitro* results indicate that α1,2-fucosidases are necessary for the utilization of fucosylated glycans (**Figure 6C**), and murine colonic mucins display this terminal epitope^11^, we hypothesized that these enzymes would be necessary to access *O*-glycans *in vivo*. The mutant lacking five α1,2-fucosidases (Δ*bt3173 +* Δ*bt4682 +* Δ*bt1777 +* Δ*bt3155 +* Δ*bt3665*) displayed no defect when competed with the WT strain (**Figure 7D**). However, in support of this hypothesis, when BT1842 was deleted in combination with the previous mutant, this mutant lacking all six α1,2 fucosidase enzymes (Δ*6*× *α1,2-Fuc*) displayed a competitive defect versus WT (**Figure 7D**). Thus, while there are functional redundancies in the linkage specificities of *B. theta* fucosidases, cumulative loss of these enzymes results in impaired utilization of *O*-glycans and *in vivo* colonization defects of *B. theta.* Collectively, these results indicate that specific enzymes targeting terminal epitopes on mucin *O*-glycans have a key role in *B. theta* fitness *in vivo*. This finding could represent a change in the current paradigm of utilization of complex glycans by PUL systems, in which endo enzymes are often key first steps. Here, we showed that the utilization of *O*-glycans also requires multiple key exo-acting enzymes that target the terminal epitopes capping these host glycans.

## Discussion

The ability of *B. theta* to express a broad suite of enzymes in response to signals contained in mucin *O*-glycans is critical for its growth on mucin-derived substrates and *in vivo* colonization^8,14,28^. Here, we further demonstrate the importance of this ability by comparing transcriptional responses of mucin PULs and other gene clusters containing potential mucin-degrading enzymes identified through their variable responses to mucin *O*-glycans from different regions of the gut or related substrates like KS. These data, combined with subsequent biochemical characterization of many of these enzymes, allowed us to identify a key set of GHs and sulfatases collectively capable of fully degrading mucin *O-*glycans.

Several studies have characterized individual or small sets of mucin-degrading CAZymes^18,20,61^. A recent study of *A. muciniphila* characterized 65 different enzymes that can cleave all of the linkages in *O*-glycans^22^. The work presented here adds a second bacterium that has a complete enzyme repertoire for degradation of *O*-glycans and to our knowledge is the first report of a complete glycan degradation pathway of colonic *O*-glycans. Of note, we characterized the activity of 33 different enzymes on complex *O*-glycans and we showed that the combined sequential activity of only 13 enzymes leads to the degradation of almost all porcine colonic *O*-glycans. Additionally, our discovery of three novel GH18 enzymes that are endo-acting on *O*-glycans is the first report of this activity inside this family. These endo-acting enzymes are likely presented at the cell surface where their activity is required to cleave long *O*-glycans into smaller oligosaccharides that can be imported into the periplasm. Previous studies have shown that cell surface endo-acting enzymes have a key role in utilization of complex glycans by *B. theta*^55,59^. Therefore, it is expected that endo-acting enzymes are critical for initiating mucin *O-*glycan degradation by *B. theta.* However, the mutant lacking all four characterized endo-acting enzymes does not exhibit a lack of growth or fitness defect. *In vitro*, *B. theta* grows best on *O*-glycans that have been chemically released from the mucin polypeptide compared to intact mucin^17^. Thus, it is possible that its endo-acting *O*-glycan-degrading enzymes are not needed during growth on these released *O*-glycans or may exist to cleave longer fragments that are less abundant on the mucins tested. It is also likely that *B. theta* encodes additional endo-enzymes. However, future studies are needed to identify such enzymes and assess their contributions to *in vitro* and *in vivo* mucin utilization phenotypes. Nevertheless, when *B. theta* is monoassociated with germ-free mice fed a low fiber diet, it is alone capable of causing thinning of the colonic mucus layer, indicating that it has sufficient enzymatic potential to elicit changes in the structure of mucin *in vivo*^27^.

This study, along with other recent studies, highlights interesting similarities and differences between the abilities of *B. theta* and *A. muciniphila* to degrade mucins. Notably, *B. theta* displays a preference for growth on free mucin *O-*glycans, while *A. muciniphila* shows a preference for mucin glycoprotein^17^. Indeed, it has been previously shown that *A. muciniphila* can import mucin fragments into the periplasm^16^. This suggests that while *B. theta* imports and utilizes short oligosaccharides, possibly generated by the *endo*-acting surface enzymes or other bacteria^17^, *A. muciniphila* imports glycoprotein fragments that are degraded in the periplasm. Therefore, it is likely that the models of mucin degradation by these two bacteria involve different key enzymes. Proteases that breakdown mucin glycoprotein are likely to play a key role in generating mucin fragments utilized by *A. muciniphila,* while the utilization of *O*-glycans by *B. theta* is dependent on enzymes targeting glycans or decorating groups such as fucose, sialic acid and sulfate. Indeed, *A. muciniphila* encodes multiple mucin proteases that tolerate different glycosylation patterns indicating that this bacterium is well adapted to the utilization of glycoproteins^62^. However, *B. theta* only encodes one protease known to be active on glycoproteins decorated with GalNAc (Tn-antigen)^63^, suggesting that this bacterium favors the utilization of *O*-glycans decorating the mucin backbone. Despite these differences, both bacteria deploy multiple enzymes that act in a sequential fashion to degrade *O*-glycans. While others have shown that *A. muciniphila* enzymes also act in a sequential fashion to degrade glycan epitopes^64^, here we deploy extensive MS studies to show how different *B. theta* enzymes specifically act on different *O*-glycans. Characterization of complete mucin-degradation pathways in multiple species will continue to illuminate key shared traits, while potentially revealing unique activities or features that may dictate the ability of a species to utilize certain mucin components or exert certain interspecies or host effects *in vivo.* Future experiments are needed to identify specific glycan cues (*e.g.*, positions of features like sulfate/fucose/sialic acid, glycan length, etc.) that induce the expression of enzymes with similar activities, which may dictate their relative importance on different substrates or *in vivo* even when redundancies exist.

An additional benefit of identifying the enzyme pathway for *O*-glycan degradation in *B. theta* is its genetic tractability, which allowed us to assess the contribution of individual enzymes in mucin utilization *in vitro* and colonization fitness *in vivo.* Loss of particular *B. theta* mucin-degrading enzymes had variable effects depending on growth substrate. This has implications for the contribution of these enzymes to the utilization of mucins from different gut regions. For example, loss of functional sulfatases results in a severe growth defect on colonic mucin *O-*glycans but does not affect growth on gastric mucin *O-*glycans. In contrast, the growth defect of the mutant lacking all six known α1,2-fucosidases was more severe on gastric mucin *O-*glycans than on colonic mucin *O-*glycans. These effects reflect known patterns of distribution of sulfation (which is more abundant in the distal gut) and fucosylation (which is more abundant in the proximal gut) in humans and pigs^12^. Thus, the substrate specificity of individual *B. theta* enzymes likely influences *in vivo* phenotypes, including the regions of the gut it can successfully colonize. The differences of phenotypes between different *O*-glycan sources also highlight the relevance of utilizing colonic mucins in future microbiota studies aiming at understanding how mucins are used by gut bacteria.

Despite encoding multiple GHs with redundant catalytic activities, the loss of specific enzymes hinders the ability of *B. theta* to utilize mucins and colonize the gut. Previously a single sulfatase was identified as a key enzyme for *B. theta* to access colonic mucins^28^. In this study we show that sialidase and fucosidase activities also play a critical role in *B. theta* utilization of *O*-glycans and *in vivo* fitness. These results indicate that enzymes targeting terminal epitopes that cap *O*-glycans play a large role in enabling *B. theta* to access mucin *O*-glycans. These findings create a therapeutic opportunity to specifically block these catalytic steps *in vivo* in cases where bacterial mucin utilization has been implicated in mucus barrier disruption and disease, such as IBD and GVHD^6,8,31^.

## Limitations of the Study

This study has focused on the degradation of porcine colonic *O*-glycans; however, the use of this substrate has several limitations. Due to differences in glycosylation between species, the translation of these results to human mucin *O*-glycans is limited. While several linkages are shared between porcine and human colonic mucins *O*-glycans, these substrates also differ in core structures, backbone length and terminal epitopes, with human mucins presenting a high abundance of core 3, longer backbones and epitopes such as Sda/Cad^12,49^. Such differences in substrate structure are likely to affect enzyme activity, and it is unclear whether the key enzymes identified in this study are important for the utilization of human mucin *O*-glycans. Additionally, the differences in glycosylation between mice and humans^49^ presents a significant limitation to translating the *in vivo* results. It is therefore important that future studies focus on validating the current study using human mucins to enable the translation of these results. Although human mucins can be obtained from human biopsies, this source does not yield enough substrate to validate all the results from this study.

In this study, we focused on the utilization of released mucin *O*-glycans, however, this substrate does not fully represent the complexity of the densely glycosylated mucin found in the gut. It was previously shown that *B. theta* does not grow *in vitro* on solubilized porcine colonic mucins^17^, however, this bacterium is able to colonize germ-free mice and affect the mucus barrier when forced to utilize host-glycans^27^. This indicates that, in the *in vivo* context, *B. theta* is likely able to utilize *O*-glycans attached to the mucin backbone. Further studies are required to determine if *B. theta* enzymes are active on intact and densely glycosylated colonic mucins and to clarify how this bacterium utilizes this complex substrate.

## Methods

### Cloning and recombinant protein production

DNA fragments encoding the predicted enzymes were amplified by PCR using appropriate primers and cloned into pET28 (NheI/ XhoI restriction sites) or pETite (Expresso T7 Cloning and Expression System, Lucigen) in frame with a N- or C-terminal His_6_ tag (**Table S13A**). For recombinant protein expression, the plasmids were transformed in *Escherichia coli* strain TUNER and cultures were grown at 37 °C in LB broth containing 50 μg/mL kanamycin to mid-exponential phase. The cultures were cooled down at 4 °C, induced with 0.2 mM isopropyl-β-D-1-thiogalactopyranoside and cultured for 16 h at 16 °C and 180 rpm. The recombinant proteins were purified from cell-free extracts by immobilized metal ion affinity chromatography as described previously^59^. The size and purity of purified proteins was checked by SDS-PAGE and the protein were dialyzed for 16h at 4 °C in 10 mM sodium phosphate, 150 mM NaCl, pH 7.5 or 10 mM MES, 150 mM NaCl, pH 7.5. Protein concentrations were determined by measuring the absorbance at 280 nm and using the molar extinction coefficient.

### Recombinant enzyme assays

The activity of recombinant enzymes against different saccharides was determined by incubating for 16 h at 37 °C at standard conditions of 1 μM of enzyme and substrates in 10 mM sodium phosphate, pH 7.0 or, in cases of sulfatases being added in the reaction, 10 mM MES, pH 6.5, with 5 mM CaCl_2_. The substrate concentration was 1 mM of di or oligosaccharides, 5 mg/mL of KS or cMO and 10 mg/mL of gMO or BSM, unless stated otherwise. Kinetics for the recombinant enzymes was determined as described previously^59^ whereas the assays for galactosidases and fucosidases was performed using the detection kits from Megazyme K-ARGA and K-FUCOSE, respectively. The activity of GH20 enzymes on pNP-β-*N*-acetylglucosamine was monitored at absorbance 405 nm. A single substrate concentration was used to calculate catalytic efficiency (k_cat_/K_M_) following the equation V_0_ = (k_cat_/K_M_)[S][E] after confirming that the substrate concentration is considerably below the K_M_ by halving and doubling the concentration and observing an appropriate increase or decrease in rate. The activity of BT3050 on β-Neu5Ac-α2,3-6S-LacNAc-pNP was monitored by measuring the absorbance at 405 nm. Initial reaction rates were determined at five substrate concentrations well below the K_M_. By plotting v_0_ against substrate concentration, a linear fit was applied, and the slope obtained was used to calculate k_cat_/K_M_. All enzymatic assays were performed in at least triplicate

### Source of purified oligosaccharides and mucin *O*-glycans

Porcine and colonic *O*-glycans were purified as previously described from porcine gastric mucins type III (Sigma) or from pig distal colon and rectum, respectively^28^. Keratan sulfate was purified from bovine cartilage powder (MP Biomedicals). Briefly, the cartilage powder was resuspended in five times (volume to weight) of 1 mM disodium EDTA, 5 mM cysteine HCl, 0.2M NaOAc, pH 6.5 and papain (10U/g of powder). The solution was incubated 24 h at 65 °C, 130 rpm with fresh cysteine and papain (20U/g of powder) being added at 12 h of incubation. The enzyme was inactivated by boiling for 5 min and the precipitated protein was removed by centrifugation at 7500 rpm for 10 min. The chondroitin and dermatan sulfate were precipitated from the supernatant by slowly adding 1.5 volumes of ethanol (under strong stirring) and removed by centrifugation as before. The peptide-keratan sulfate was recovered from the supernatant by ethanol precipitation (1.5 volume) for 16 h at 4 °C followed by centrifugation at 7500 rpm, 4 °C for 10 min. The purified keratan sulfate was resuspended and dialyzed (10 kDa cut-off) extensively against water before being freeze-dried.

PSGL-1/mIgG2b fusion proteins were expressed in CHO cell pools transiently transfected with cDNAs encoding specific *O*-glycan core enzymes and secreted fusion proteins were collected from culture supernatants and purified using Protein G affinity chromatography as previously described^65^.

The sialylated O-glycans (Neu5Ac-α2,3-Gal-β1,4-6SGlcNAc-β1,3-Gal-β1,3-GalNAc-α-C_2_- N_3_ and Neu5Ac-α2,6-Gal-β1,4-6SGlcNAc-β1,3-Gal-β1,3-GalNAc-α-C_2_-N_3_) were prepared as previously described^66^. Core 2 and 4 were synthetized as previously described^67–69^. The remaining oligosaccharides were from Dextra Laboratories, Carbosynth, Elicityl, Megazyme and Sigma.

### Thin layer chromatography (TLC) and high-performance anion exchange chromatography with pulsed amperometric detection (HPAEC-PAD)

For visualization of enzymatic reactions by TLC, 3 μL samples were spotted twice onto TLC Silica gel 60 plates (Millipore) and resolve in butanol:acetic acid:water (2:1:1). Plates were dried and sugars were visualized using diphenylamine developer (2 g diphenylamine, 100 mL ethyl acetate, 2 mL aniline, 10 mL H_3_PO_4_, 1 mL HCl) and heating over a flame or incubation at 100 °C for 20 minutes. When necessary, HPAEC-PAD was used to confirm the enzymatic activity by using appropriated standards. Briefly, the substrates and reaction products were bound to a Dionex CarboPac P100 column and eluted (flow rate 1.0 mL min^−1^) with an initial isocratic flow of 10 mM NaOH for 20 min, followed by a gradient of 10–100 mM NaOH for 20 min, and an isocratic flow of 100 mM NaOH and 500 mM sodium acetate. For reactions with endo-acting GH18 enzymes, the elution was performed with an initial isocratic flow of 100 mM NaOH for 20 min, followed by a gradient of 0–100 mM sodium acetate for 20 min, and an isocratic flow of 500 mM sodium acetate in100 mM NaOH. The quantification of fucose released by different fucosidases was calculated using a standard curve. Briefly, the areas under the curve of the peak for fucose standards of known concentration were used to generate a standard curve that was used for the quantification of released fucose in different enzymatic reactions.

### Liquid chromatography with electrospray ionization tandem mass spectrometry

To release *O*-glycans from glycoprotein (BSM and PSGL-1/mIgG2b), proteins were blotted to PVDF membrane (Bio-Rad). After blotting, proteins bands were visualized by direct blue 71 staining (0.008% in 10% acetic acid and 40% ethanol, v/v). Bands were excised and subjected to reductive β-elimination. In brief, membrane strips were incubated with 20 μL of 0.5 M NaBH_4_ and 50 mM NaOH for16 h at 50◦C. Reactions were quenched with 1 μL of glacial acetic acid, and samples were desalted and dried as previously described^70^. For released oligosaccharides (cMO, gMO, and KS), the samples were cleaned up with porous graphitized carbon (PGC) solid-phase extraction (Thermo Scientific). After loading onto PGC cartridges, the oligosaccharides were washed and eluted with SEP elution solution. The eluates were then dried and resuspended in water prior to analysis. Released *O*-glycans were analyzed by liquid-chromatography-mass spectrometry (LC-MS) using a 10 cm × 250 μm I.D. column, prepared in-house, containing 5 μm porous graphitized carbon (PGC) particles (Thermo Scientific). Glycans were eluted using a linear gradient from 0–40% acetonitrile in 10 mm ammonium bicarbonate over 40 min at a flow rate of 6 μL/min. The eluted *O*-glycans were detected using an LTQ mass spectrometer (Thermo Scientific) in negative-ion mode with an electrospray voltage of 3.5 kV, capillary voltage of −33.0 V and capillary temperature of 300 °C. Air was used as a sheath gas and mass ranges were defined depending on the specific structure to be analyzed. The data were processed using Xcalibur software (version 2.0.7, Thermo Scientific). Glycans were annotated from their MS/MS spectra manually and validated by available structures stored in UniCarb-DB database (2025 version)^71^. For structural annotation, some assumptions were used in this study: monosaccharides in the reducing end were assumed as GalNAcol; GalNAc was used for HexNAc when identified in blood group A and core 5 *O*-glycans, otherwise HexNAc was assumed to be GlcNAc; hexose was interpreted as Gal residues. *O*-glycans with linear cores (core 1, 3, and 5) were distinguished from branched cores (core 2 and 4) based on the presence of [M - H]^−^ − 223 and [M - H]^−^ - C_3_H_8_O_4_ (or 108 au) in MS/MS of structures with linear core^72^.

### Anaerobic bacterial culture and genetic manipulation

*B. theta* VPI-5482 *tdk^-/-^* was the parent strain utilized as wild-type in this study and the strain from which additional mutant strains were generated. Mutant strain gene deletions were generated using counter-selectable allelic exchange as previously described^73^ and the primers are listed on **Table S13B**. *B. theta* was cultured anaerobically (10% H2, 5% CO2, 85% N2; Coy Laboratory Products) at 37°C in TYG medium or minimal medium (MM)^14^ containing the appropriate carbon source. Briefly, overnight cultures were washed and diluted 1:40 and resuspended in 2X concentration MM^73^ and 100 μl was added to the substrates (2X concentration, 100 ul) in 96 well plates (Costar). The increase in absorbance at 600 nm was collected to measure growth (Biotek). Growth curves were normalized by subtracting the average absorbance values of the first three readings from all growth values to account for absorbance of growth substrates. The data collected was analyzed in GraphPad Prism v10. All growth curves presented are the averages and s.e.m. of three technical replicates, except for growth curves on cMO and Tetra-LacNAc where two technical replicates were used due to the limited amount of these substrates.

### RNA extraction, sequencing and data processing

We performed RNA-sequencing (RNA-seq) analysis to measure changes in *B. theta* gene expression during growth in MM containing porcine colonic mucin *O*-glycans (cMO, 5 mg/ml), porcine gastric *O*-glycans (gMO, 10 mg/ml), or type II keratan sulfate (KS, 5 mg/ml). Tetra *N*-acetyllactosamine (Tetra-LacNAc, 5 mg/ml) was tested to determine the stimulatory effects of an unsulfated octasaccharide that resembles the backbone structure of many *O*- glycans but was not used to directly identify candidate mucin PULs (*i.e*., if a PUL gene was only upregulated during growth in LacNAc_4_ and none of the other three substrates, it was not included in **Figure 1B**, although expression changes for all genes are shown in **Table S1A**). Growth in MM-glucose (5mg/ml) was use as a reference condition for differential expression analysis. For a gene to be considered differentially regulated in one or more conditions its transcription level needed to differ by ≥10-fold relative to glucose. RNA-seq data were gathered on bulk RNA that was extracted as previously reported^8^, depleted using Ribozero reagents (Epicentre, Madison, WI) and sequenced on the Illumina HiSeq platform with 50bp reads. Reads were mapped to the *B. theta* genome using Arraystar software (Lasergene, Madison, WI) using reads per kilobase million (RPKM) normalization as previously reported^17^.

To reinforce the potential involvement of candidate *O*-glycan PULs, we added back the additional PUL genes present in each locus even if they were not expressed ≥10-fold, which reduces contributions of spurious signals from one or a few genes in a PUL and also provides a better estimate of expression intensity by averaging measurements from multiple genes^14^. More complex PULs were analyzed by separating out individual operons or genes based on manual inspection of coordinated expression behavior and transcription start sites reported in ThetaBase^74^. The expression levels of genes in each PUL or individual PUL operon were then averaged and all PULs with at least one operon, or individually expressed genes in a few rare cases, were retained for further analysis (**Table S1C**). Because the cMO and KS substrates appeared to be contaminated with small amounts of other polysaccharides (cMO likely from incomplete removal of dietary fiber/feces from pig colons; KS from incomplete separation of KS from chondroitin sulfate in cartilage), PULs that have been previously shown to respond to purified dietary fibers (*e.g*., arabinan, arabinogalactan, fructans) or chondroitin sulfate were noted in **Table S1B** and removed from further consideration as candidate *O*-glycan PULs. A more difficult distinction is potential overlap between PULs that target *N*- and *O*-linked glycans because several known PULs with enzymes specific for *N*-linked glycans are also upregulated in response to growth on our *O*- glycan preparations either because our preparations contain contaminating *N*-glycans, which are also found in mucin at lower levels, or because these PULs are expressed in response to sub-motifs that are present in both glycan types. While this overlap cannot be completely unraveled due to the heterogeneity of both glycan types, we excluded PULs that were previously shown to be expressed during growth in α1-acid-glycoprotein, which is only known to contain *N*-glycans^44^. Therefore, PULs that are both induced by this substrate and encode enzymes (mostly GH18s that do not cluster with the GH18 endo-*O*-glycanases described in this study) that degrade *N*-glycan specific linkages were omitted from further consideration. The final list of PULs that are candidates for *O*-glycan foraging are listed in **Table S1D**.

### Immunolabeling of *B. theta* cell surface proteins

Polyclonal antibodies were raised in rabbits immunized with individual recombinant enzymes BT1629 and BT1632 (Cocalico Biologicals). Antibodies were purified from serum by incubating serum 1:1 by volume with a culture of the mutant deletion of PUL *bt1623-1636*, pelleting cells, and collecting the supernatant. To localize enzymes, *B. theta* was grown overnight in TYG and back-diluted 1:80 in MM with keratan sulfate (5 mg/ml), and grown for 6.5 h. Bacteria were fixed in formalin for 1. 5h at room temperature, pelleted, and washed twice with PBS. Cells were pelleted and resuspended in 1 mL of blocking solution (2% goat serum, 0.02% NaN_3_ in PBS) overnight at 4 °C. All further incubations of antibodies were done in blocking solution. Cells were pelleted and stained with 0.5 mL of purified antibody solution (1:500) for 2 h at room temperature. After washing cells 3 times in PBS, and the cell pellet was resuspended in 0.4 mL of goat anti-rabbit Alexa Fluor 488 (1:500) and incubated for 1 h at room temperature. Cells were washed three times in PBS and resuspended in 20 μL PBS and 1 drop of Prolong Gold Antifade. Samples were stored covered at 4 °C until images were acquired with Zeiss Apotome microscope.

### Detection of *B. theta* surface enzyme activity

*B. theta* wild-type and the deletion mutant of two fucosidases (Δ*bt3173* + Δ*bt4682*) were cultured until mid-log phase in 5 ml of MM supplemented with porcine gastric *O*-glycans (gMO, 10 mg/ml). The bacteria were isolated by centrifugation at 4500 rpm for 5 min at room temperature, the pellets were washed once with PBS and resuspended into 3 ml of PBS. 1 ml of the bacterial suspension was kept on ice (intact cells) and the remaining 2 ml were sonicated to lyse the bacteria and release intracellular enzymes (sonicated cells). Intact and sonicated cells were incubated at 37 °C with equal amounts (1:1) of gMO (final concentration 10 mg/ml in PBS). A control reaction was set up where gMO was incubated under the same conditions but only with PBS. Samples were collected at different time points and immediately boiled 5 min to stop enzymatic activity. To determine gMO degradation in the different samples, the reactions were analyzed by TLC and HPAEC.

### Gnotobiotic mouse experiments

Mouse experiments were approved by the University Committee on Use and Care of Animals at the University of Michigan (NIH Office of Laboratory Animal Welfare number A3114– 01). Germ-free Swiss Webster mice male and female (6-8 weeks old) were weaned onto a fiber deficient diet (TD130343, Envigo) for seven days prior to oral gavage and this diet was maintained throughout the experiment. At day 0, mice were gavaged with equal amounts of *B. theta* wild-type and mutant. Fecal samples were collected at different time-points up to 42 days, with the exception of one experiment where samples were collected up to 77 days. The extraction of bacterial genomic DNA and the quantification of relative abundance by qPCR was carried out as previously described^28^. For samples with low abundance of at least one tagged strain by qPCR (below the lowest concentration on the standard curve), samples were repeated after adding 20 ng per well and results were reported for these second analyses.

### Bioinformatics and visualization of structures

All protein sequences were retrieved from CAZy^24^ or KEGG databases^75^. The proteins predicted signal peptides (SP) were identified using the LipoP1.0 server^76^. All protein sequences or protein domains were aligned with MUSCLE^77^ and the percentage sequence identity of the catalytic domains and putative binding domains (F5/8 type C) retrieved. To generate the GH18 phylogenetic tree the protein sequences of previously characterized members and the sequences of all GH18 from *Bacteroides* type strains were downloaded from CAZy database^24^. Sequences were aligned as previously described and the tree was generated in MEGA 11^78^. Briefly, the evolutionary analysis was inferred by using the Maximum Likelihood method and JTT matrix-based model^79^ and the bootstrap consensus tree inferred from 50 replicates^80^. The structures of BT1627 and BT3178 were generated with AlphaFold3^81^ and the superimposition was performed in WinCoot using the Secondary Structure Matching^82^. Protein structures were visualized using PyMOL.

### Quantification and statistical analysis

Statistical analyses were carried out in GraphPad Prism v10.5.0 at all timepoints with at least three samples, and two-tailed, paired *t*-tests.

## Acknowledgements

Funding was provided to ASL by Swedish Research Council Grant (2021-01409), Swedish Society for Medical Research (Svenska Sallskapet for Medicinsk Forskning, Grant S21-0026), Sahlgrenska Academy International Starting Grant (GU2021/1070), and Jeanssons Foundation Grant. ECM, SRS, ASL and CJ were supported by funds from the US National Institutes of Health (R01 DK125445 to ECM). CRC and RH gratefully acknowledge funding from the Swiss National Science Foundation (CRSK-3_196773 & 320030-231409 to RH), Innosuisse (104.462 IP-LS to RH), and the University of Basel. GR was funded by the Estonian Research Council (PUTJD1227). SRS was supported by funds from the NIH T32 Cellular Biotechnology Training Program (T32 GM145304). JH was funded by grant ALFGBG-1006863 from the Swedish state under the agreement between the Swedish government and the country councils. ASL was previously supported by European Union’s Horizon 2020 research and innovation program under Marie Skłodowska-Curie grant agreement no.748336. We thank SciLifeLab and BioMS funded by the Swedish research council for providing financial support to the Proteomics Core Facility, Sahlgrenska Academy.

## Author contributions

E.M. and A.S.L designed the experiments and wrote the manuscript. S.R.S. and A.S.L cloned all the putative enzymes. S.R.S., R.B, M.J.G-B., M.N., N.D.S. and A.S.L performed protein expression and purification. S.R.S, N.A.P and E.C.M. and A.S.L performed the extraction of mucins and keratan sulfate. S.R.S, R.B., M.J.G-B., G.V.P., M.N., N.D.S. and A.S.L conducted quantitative and qualitative enzyme assays. C.J. performed mass spectrometry analyses. S.R.S., A.S.L and E.M. performed the RNA-sequencing analysis. R.B., S.G.W, J.L., J.H., C.R.C and R.H. performed the synthesis of glycans. S.R.S., G.R., A.S.L. generated the deletion strains and performed bacterial growth experiments. S.R.S., Q.Y and A.S.L carried out the gnotobiotic mouse experiments. S.R.S. carried out the cellular localization by immunolabeling. All authors read and approved the manuscript.

## Data availability

The full RNA-Seq data provided in **Table S1** was submitted to https://www.ncbi.nlm.nih.gov/geo/ with the accession number pending. Glycomics raw files were uploaded to Glycopost (https://glycopost.glycosmos.org/preview/812254461690db0cc5b701, password 5613)

---

**Figure S1. *B. theta* growth on different host glycans upregulates different PULs**

(A) Growth of *B. theta* wild-type (*Bt* WT) on gMO (10 mg/mL), cMO (5 mg/mL), KS (10 mg/mL) or Tetra-LacNAc (5mg/mL). Lines represent the mean of biological replicates (*n* = 3 gMO and KS; n =2 cMO and Tetra-LAcNAc; error bars, SEM). Please note that gMO, cMO and KS curves are also shown for the WT growth in Figure 6A.

(B) Venn diagram of PUL genes upregulated during growth on gMO, cMO, and KS.

(C) Heatmap showing the average fold-change (relative to MM-glucose) of PULs upregulated during growth on gMO, cMO, and KS.

cMO, porcine colonic mucin *O*-glycans; gMO, porcine gasric mucin *O*-glycans; KS, keratan sulfate; Tetra-LacNAc, tetra-*N*-acetyllactosamine.

**Figure S2. *B. theta* growth on different host glycans upregulates different PULs**

Schematic representation of the molecular architecture of the characterized enzymes showing their domains, amino acid (aa) length, and predicted signal peptides (SP). SP I denotes a signal peptide recognized by signal peptidase I, while SP II denotes a lipoprotein signal peptide cleaved by signal peptidase II. For each family, the percentage sequence identity of the catalytic domains and putative binding domains (F5/8 type C) is shown on the right. GHXX, glycoside hydrolase family XX; CBMXX, carbohydrate-binding module family XX; UNK, domain of unknown function.

**Figure S3. GH18 family contains endo-enzymes active on host glycans**

(A) Phylogenetic tree of GH18 family members showing previously characterized enzymes (*) and *B. theta* GH18 enzymes. The color code indicates the predicted activity based on phylogenetic similarity to previously characterized chitinases (orange) and endo-*N*-acetylglucosamidases (green). Two branches (blue and purple) do not contain any previously characterized enzymes.

(B) Relative abundance of different oligosaccharides detected by mass spectrometry in reactions with BT1632 or BT3050 and various glycans. The detected glycans are color-coded, and the numbers and structures for the specific *m/z* values are shown below. The “no enzyme” control corresponds to reactions carried out under the same conditions without enzyme.

(C) Digestion of the reaction products generated by endo-acting enzymes BT1632 and BT3050 on KS, analyzed by thin-layer chromatography. The reactions were incubated (+) with 1 µM sialidase (BT0455), 6S-Gal sulfatase (BT1624), and β1,4-galactosidase (BT1626) for 16 h at 37 °C. The main products of the endo-acting enzymes are labeled with numbers (1 to 3) and dashed lines in the corresponding panels. Standards and their respective labels are shown on the left side. KS, keratan sulfate; Gal, galactose.

**Figure S4. Activity of enzymes cleaving terminal epitopes**

(A to C) Analysis of the reaction products from fucosidase activity against terminal fucosylated oligosaccharides by TLC and HPAEC-PAD detection.

(D) Glycans detected by mass spectrometry in cMO before (control) and after fucosidase treatment. Relative abundance for specific *m/z* glycans (left panel) and glycans with terminal fucosylated linkages (right panel) (n = 3; error bars = SEM).

(E) Activity of GH110 against oligosaccharides detected by TLC. For blood group B type I and II, the substrate was incubated with GH110 alone or in combination with different fucosidases (GH29 and GH95).

(F) Reaction products of GH109 enzymes against different substrates, detected by TLC and HPAEC-PAD.

(G) TLC detection of GlcNAc released by GH89 incubated with gMO

(H) Reaction products of sialidase incubated with different substrates, detected by TLC. * indicates detection of sialic acid.

(I) Relative abundance of oligosaccharides with or without terminal sialic acid epitopes before (control) and after incubation with sialidase (left) and structures for the specific glycans not detected after enzymatic treatment (right)

TLC and HPAEC-PAD standards are shown on the left or top, respectively. Where possible, substrates and reaction products are identified. When applicable, the locus tag of each enzyme is followed by its GH family in superscript. cMO, porcine colonic mucin *O*-glycans; gMO, porcine gastric mucin *O*-glycans; PGM, porcine gastric mucin; BSM, bovine submaxillary mucin.

**Figure S5. Activity of *B. theta* GH2 and GH20 enzymes**

(A) Analysis by TLC and HPAEC-PAD analysis of reaction products generated by GH2 activity against oligosaccharides. All reactions set in standard conditions with exception of core 2 reactions that were set either with 1 or 0,01μM of enzyme.

(B) Activity of GH2 enzymes against sulfated glycans detected by TLC. Sulfated 6S-LacNAc and 6S-LNB were generated by incubating 6-Lewis x and 6-Lewis a with 1 μM of fucosidase (BT1625), respectively. KS oligosaccharides were generated upon treatment with 1 μM of BT1632.

(C) Glycans detected by mass spectrometry in BSM (bovine submaxillary mucin) before (control) and after galactosidase treatment (left). Relative abundance and corresponding structures for specific *m/z* values are shown on the right. Ions with an underscore followed by a number represent glycans with the same mass but different structures.

(D) Relative abundances and corresponding structures for specific *m/z* glycans detected on P-selectin glycoprotein ligand-1 (PSGL1) after incubation with GH2 enzymes. PSGL1 was incubated with 1 μM sialidase (BT0455) to enrich for core structures.

(E) TLC analysis of reaction products from GH20 enzymes. For all reactions, LNnT was incubated with 1 μM of galactosidase (BT1626) to generate a trisaccharide with a non-reducing GlcNAc.

(F) TLC and HPAEC-PAD analysis of reaction products of Tetra-LacNAc incubated with 1 μM of galactosidase (BT1626) in presence or absence of GH20 enzymes.

(G) Surface representation of BT1627 and BT3178 structures predicted with AlphaFold3. The structures were overlaid with Amuc_3136 (PDB 6JQF) and GlcNAc is shown in active site (purple). Residues not conserved between both proteins are shown as sticks and labeled in red and underlined. The catalytic residues are shown as sticks and labeled in black.

(H) TLC analysis of reaction products of GH20 enzymes against core 2 (1 mM) pre-treated with 1 μM of galactosidase (BT1626) for 9 h and against core 4 (0.5 mM) pre-treated with 50 nM of BT3868 for 30 min, to generate the disaccharide GlcNAc-β1,6-GalNAc-Methyl.

(I) Reaction products of 50 nM of BT3868 against core 4 (0.5 mM) in different time-points analyzed by TLC, with relative abundances of substrate and products detected by mass spectrometry.

In all reactions, control represents reactions performed without tested enzymes under identical conditions. TLC and HPAEC-PAD standards are shown on the left or top, respectively. Where possible, substrates and reaction products are identified. When applicable, the locus tag of each enzyme is followed by its GH family in superscript. LNB, lacto-*N*-biose; LNT, lacto-*N*-tetraose; Di-LacNAc, Di-*N*-acetyl-D-lactosamine; LNFP IV, lacto*-N*-fucopentaose IV; LNnT, lacto-*N*-neotetraose; KS, keratan sulfate; GlcNAc, *N*-acetyl-D-glucosamine.

**Figure S6. Activity of *B. theta* GH2 and GH20 enzymes**

(A) Sequential degradation of Blood group A and B trisaccharides analyzed by HPAEC-PAD. Standards and reaction products are labelled at the top. The initial substrate is highlighted in grey.

(B to D) TLC analyses of the products generated during sequential degradation of fucosylated, sulfated and sialylated oligosaccharides. Enzymes included in each reaction are indicated below the TLC (+). Standards are shown on the left and labeled at the top.

(E) Schematic representation of the enzymatic activities of *B. theta* enzymes included in the enzymatic degradation of colonic *O*-glycans. Arrows indicate the specific linkages targeted by each enzyme. Enzyme mix 1 contained all the enzymes except BT4394, which was included only in enzyme mix 2. When applicable, enzyme locus tags are followed by their respective GH family in superscript.

**Figure S7. *B. theta* enzymes activities required to growth and utilization of *O*-glycans**

(A) Immunofluorescence and differential interference contrast (DIC) microscopy of wild-type *B. theta* stained with polyclonal antibody (green, Anti-BTXXXX) against BT1629 or BT1632 and DAPI for DNA (blue). Scale bar, 5 μm.

(B) TLC analysis of oligosaccharides present in the culture supernatant of *B. theta* wild-type (*Bt* WT) or gene-deletion mutants (Δ*btXXXX*) after growth on gMO for 5 days.

(C) Growth of *Bt* WT and fucosidase mutant in presence or absence of 1 μM of active fucosidases (BT3173^GH95^ and BT4682^GH95^) on gMO (10 mg/ml). Lines represent the mean of biological replicates (n = 3; error bars, SEM). The right panels show the HPAEC-PAD and TLC analysis of the oligosaccharides in culture supernatant after growth.

(D) Left panel: TLC analysis of enzymatic activity at the surface of intact cells (IC) or total enzymatic activity in the bacteria after intracellular enzymes being released by sonication (sonicated cells, SC). IC and SC were incubated aerobically at 37 °C with gMO (10 mg/ml) and samples were collected at different time-points (0h, 2h, 3h and 16h). The gMO control sample represents the substrate incubated with PBS in the absence of bacteria. Right panel: HPAEC-PAD analysis of monosaccharides detected at time 0 and after 16 h of incubation with IC or SC.

(E) Growth of *B. theta* wild-type (Bt *WT*) and gene-deletion mutants (Δ*btxxxx*) on cMO (5 mg/ml). Lines represent the mean of biological replicates (n = 3; error bars, SEM).

cMO, porcine colonic mucin *O*-glycans; gMO, porcine gastric mucin *O*-glycans;

