## Supplementary figures and images for "Degradation of mucin *O*-glycans by a human gut symbiont requires a complex enzyme repertoire and promotes colonization"

### Supplemental figures

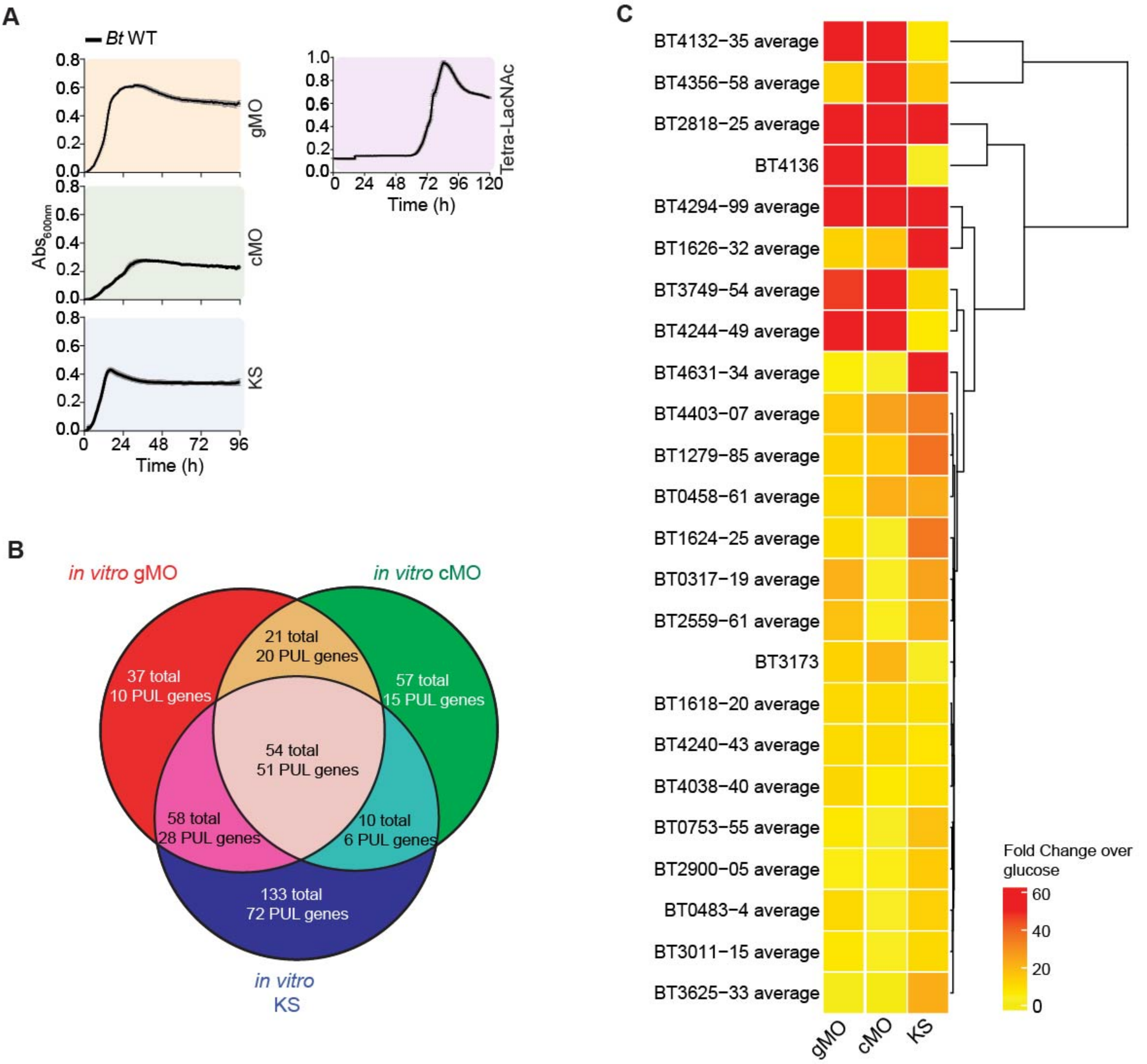

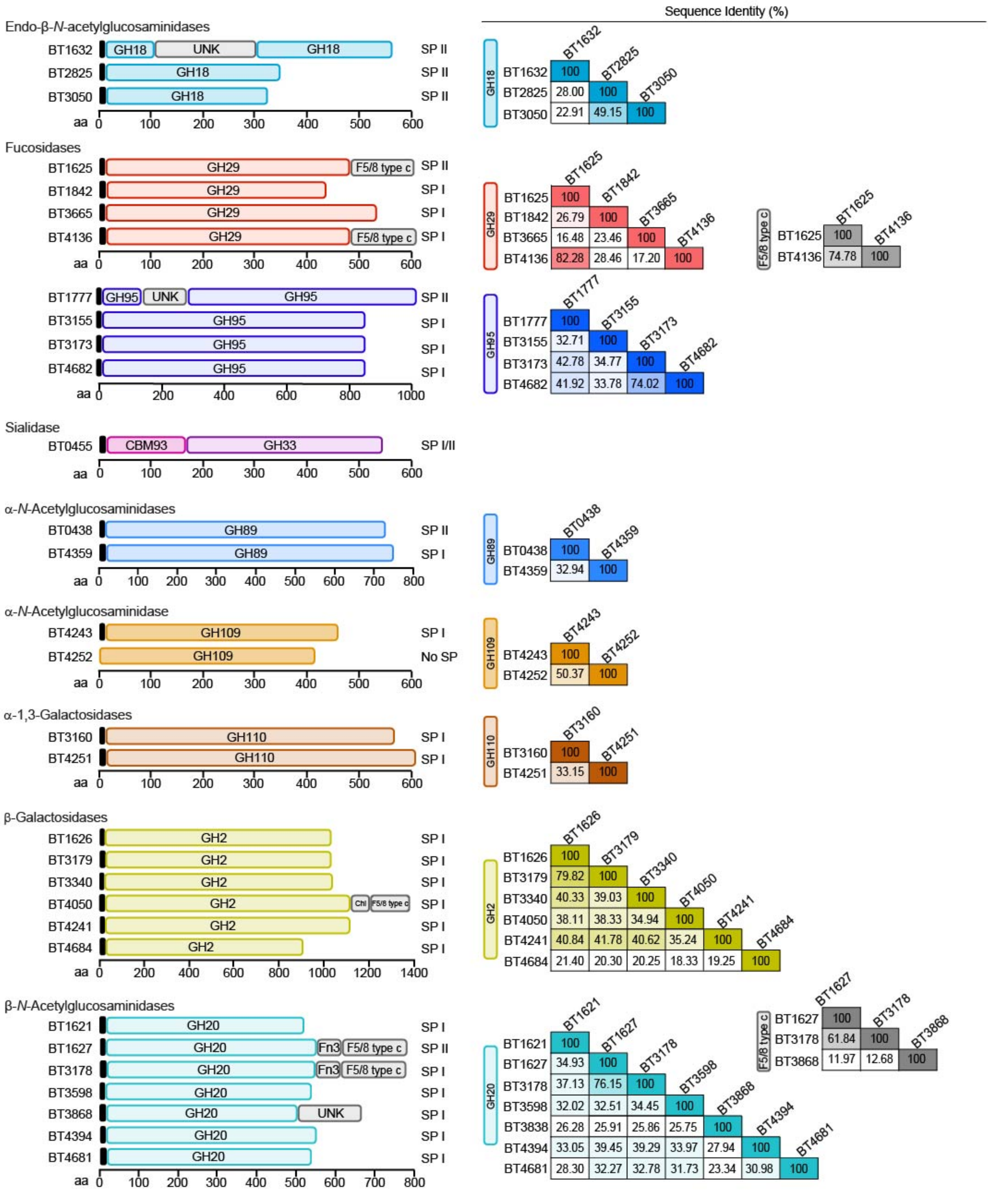

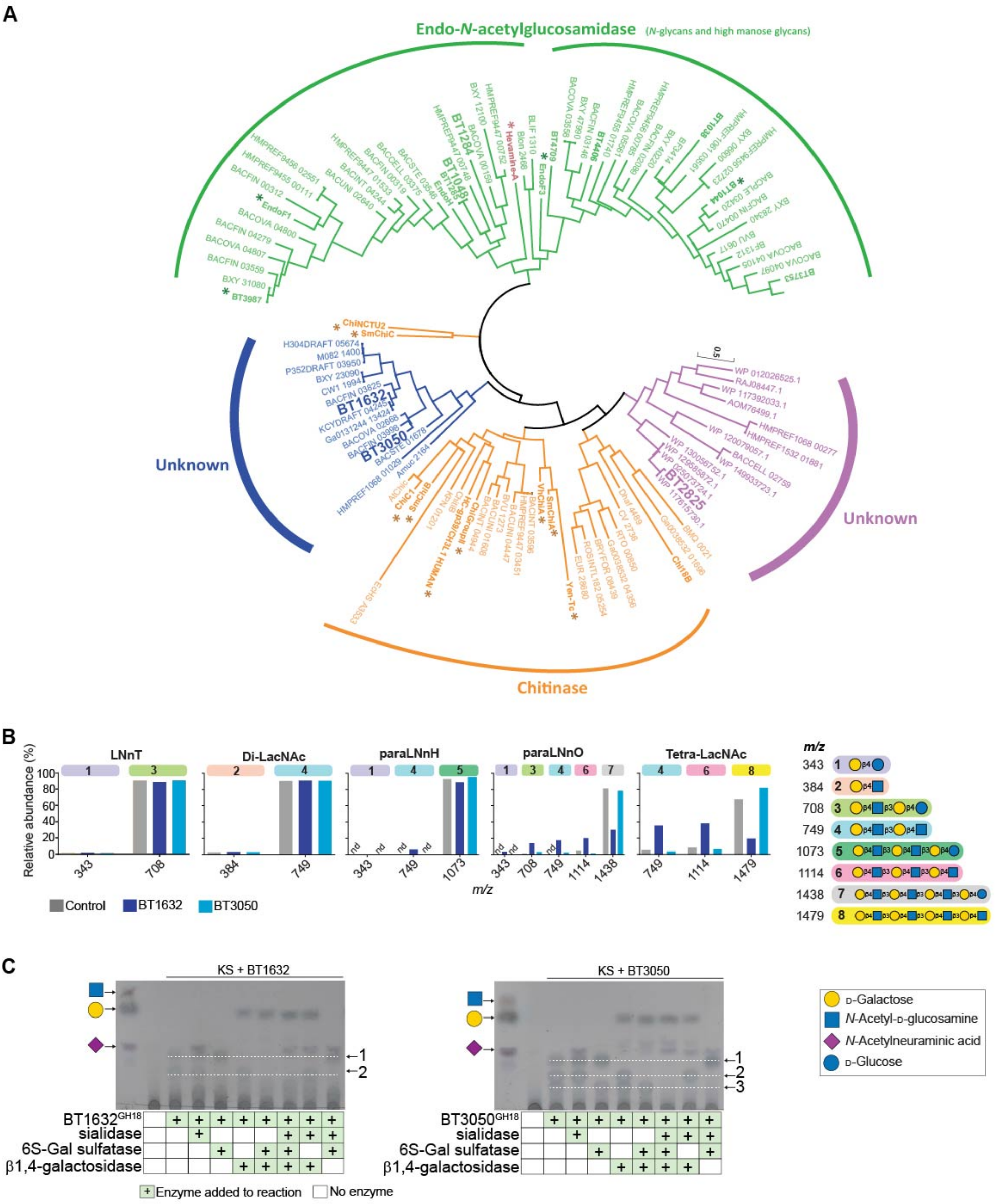

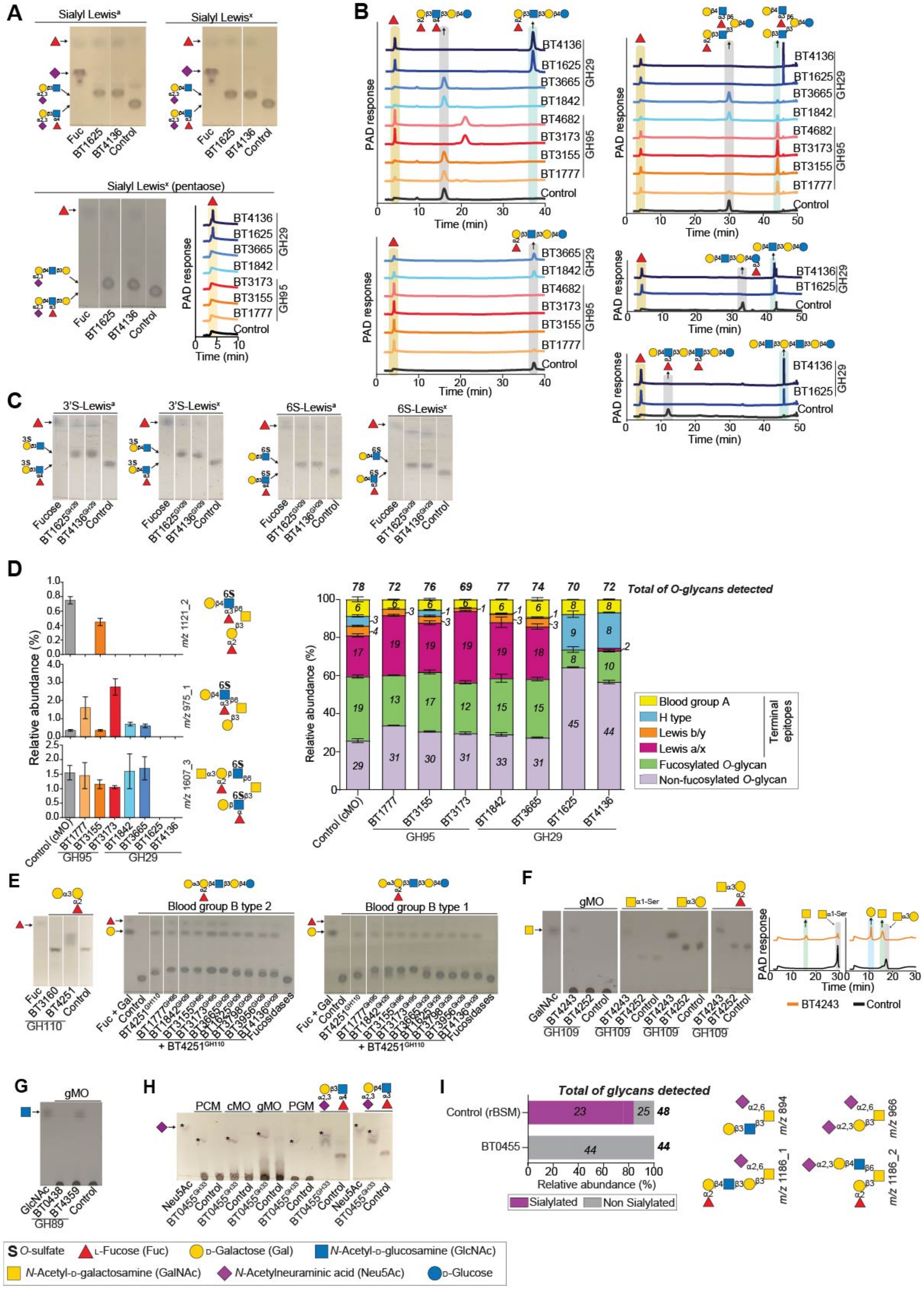

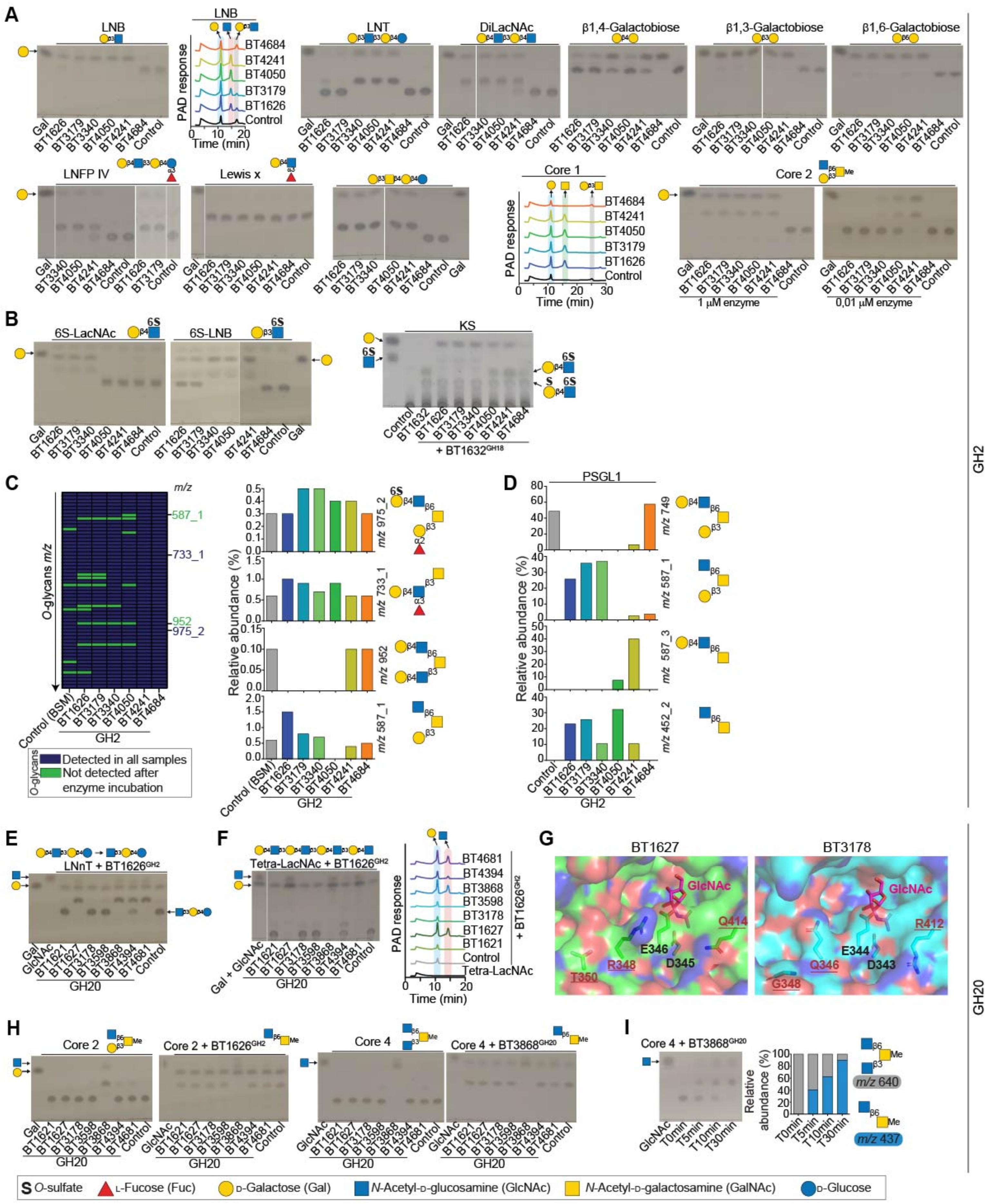

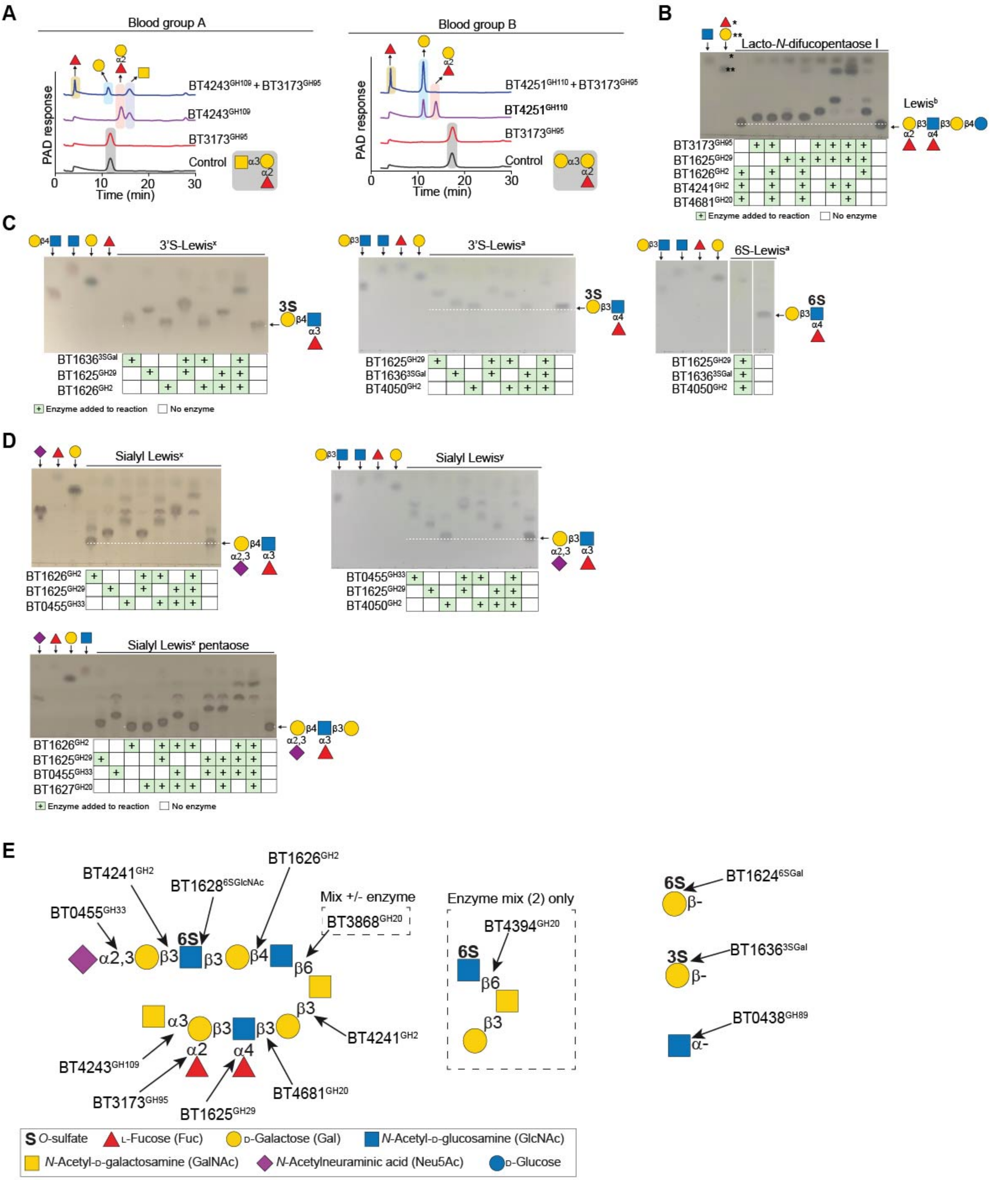

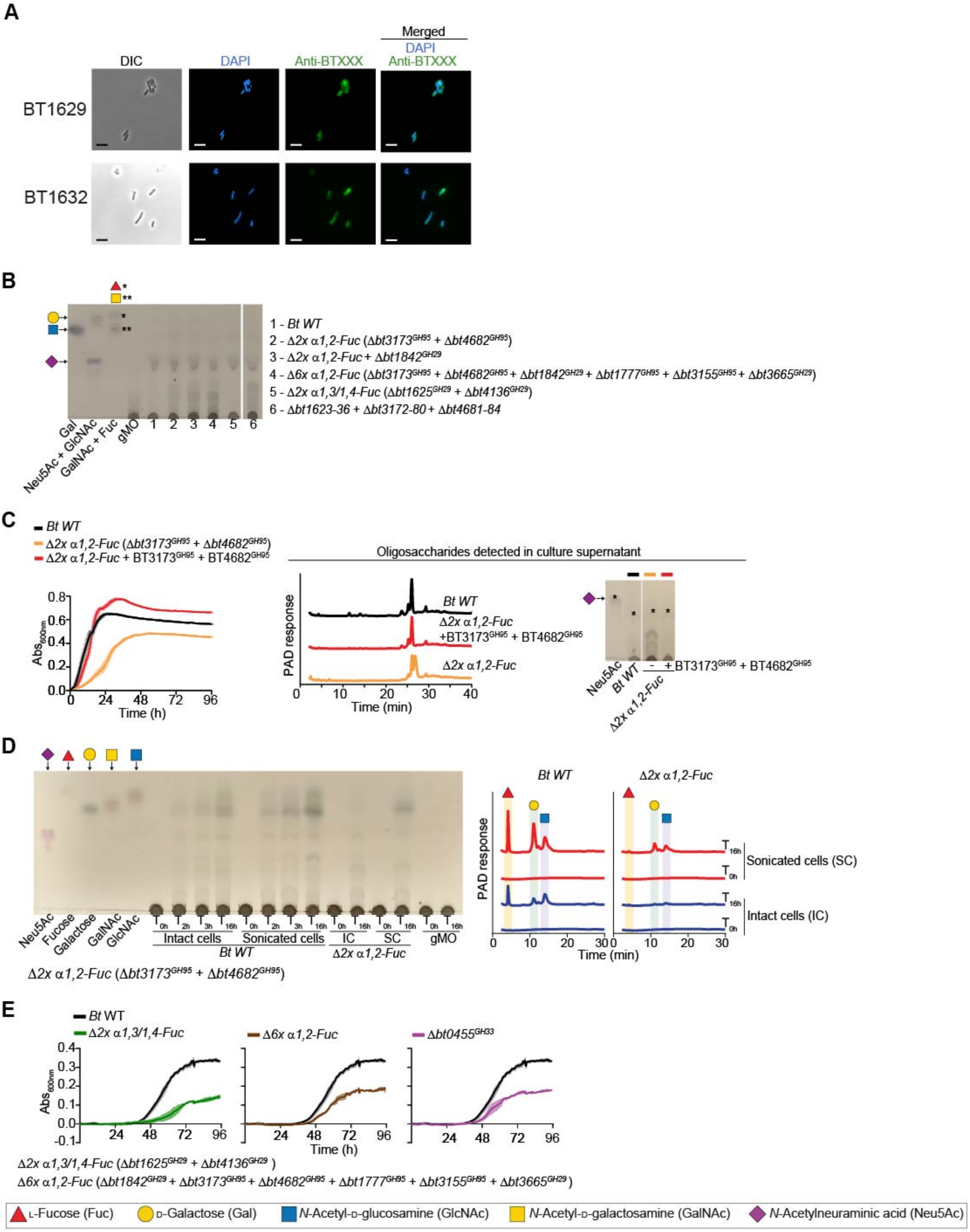
